# Developmental HCN channelopathy results in decreased neural progenitor proliferation and microcephaly in mice

**DOI:** 10.1101/2021.04.24.441237

**Authors:** Anna Katharina Schlusche, Sabine Ulrike Vay, Niklas Kleinenkuhnen, Steffi Sandke, Rafael Campos-Martin, Marta Florio, Wieland Huttner, Achim Tresch, Jochen Roeper, Maria Adele Rueger, Igor Jakovcevski, Malte Stockebrand, Dirk Isbrandt

## Abstract

The development of the cerebral cortex relies on the controlled division of neural stem and progenitor cells. The requirement for precise spatiotemporal control of proliferation and cell fate places a high demand on the cell division machinery, and defective cell division can cause microcephaly and other brain malformations. Cell-extrinsic and intrinsic factors govern the capacity of cortical progenitors to produce large numbers of neurons and glia within a short developmental time window. In particular, ion channels shape the intrinsic biophysical properties of precursor cells and neurons and control their membrane potential throughout the cell cycle. We found that hyperpolarization-activated cyclic nucleotide-gated cation (HCN)-channel subunits are expressed in mouse, rat, and human neural progenitors. Loss of HCN-channel function in rat neural stem cells impaired their proliferation by affecting the cell-cycle progression, causing G1 accumulation and dysregulation of genes associated with human microcephaly. Transgene-mediated, dominant-negative loss of HCN-channel function in the embryonic mouse telencephalon resulted in pronounced microcephaly. Together, our findings suggest a novel role for HCN-channel subunits as a part of a general mechanism influencing cortical development in mammals.

**Significance Statement:** Impaired cell cycle regulation of neural stem and progenitor cells can affect cortical development and cause microcephaly. During cell cycle progression, the cellular membrane potential changes through the activity of ion channels and tends to be more depolarized in proliferating cells. HCN channels, which mediate a depolarizing current in neurons and cardiac cells, are linked to neurodevelopmental diseases, also contribute to the control of cell-cycle progression and proliferation of neuronal precursor cells. In this study, HCN-channel deficiency during embryonic and fetal brain development resulted in marked microcephaly of mice designed to be deficient in HCN-channel function in dorsal forebrain progenitors. The findings suggest that HCN-channel subunits are part of a general mechanism influencing cortical development in mammals.

## INTRODUCTION

Cell proliferation is a tightly regulated process consisting of multifaceted and complex control mechanisms. Defective cell-cycle checkpoints, or the dysfunction of cell cycle regulators, affect the physiological or pathological fate of stem and neural progenitor cells (NPCs), and defects in these processes can cause microcephaly and other brain malformations (1). While the molecular and biochemical mechanisms of cell-cycle control are well established, the bioelectrical modulation of cell-cycle progression by ion channels is still poorly understood. Ion channel activity has been suggested to regulate proliferation through its effect on the membrane potential. Notably, it had been shown that non-proliferative cells had lower resting membrane potential than proliferative cells and that the membrane potential changed during cell-cycle progression (2, 3). Hyperpolarization of dividing tumor cells induced cell-cycle arrest (2), while strong depolarization of even post-mitotic neurons induced their proliferation (4, 5). Furthermore, a recent study showed that the resting membrane potential of mouse cortical progenitors influenced their neurogenic differentiation (6). During development, the profile of ion channels expressed by progenitor cells exhibits changes, and so do the biophysical properties they are endowed with (7–9). In particular, K^+^ channels, such as the *ether-a-go-go* gene family members EAG and ERG, have been linked to cell-cycle progression due to their cell-cycle phase-specific expression and function (for review see (3)). Together, the data, which were predominantly obtained from cultured and tumor cell lines, suggest a close relationship between membrane potential and the well-investigated cyclin-dependent pathways. Altered function of ion channels that contribute to the control of the membrane potential in neural stem and progenitor cells could, therefore, affect proliferation or differentiation and impair brain development.

Recently, several patients with early infantile epileptic encephalopathy (EIEE) were reported to have mutations in *HCN1*, which encodes a pore-forming subunit of hyperpolarization-activated cyclic nucleotide-gated cation (HCN) channels (10, 11). Two unrelated patients carried the *HCN1* missense loss-of-function mutation p.M305L and presented with epilepsy and a deceleration of head growth resulting in microcephaly (10). In the absence of overt signs of structural changes in these patients, such as altered gyration, focal dysplasia, or enlarged subarachnoidal space or ventricles in cerebral magnetic resonance tomography, we sought to investigate whether *HCN* channelopathies might affect processes of brain development that could impair brain growth. Mutations in *HCN1* are associated with a broad spectrum of developmental disorders, including intellectual disability and autism spectrum disorder, atypical Rett syndrome with microcephaly (11), generalized epilepsy, febrile seizures plus spectrum, and EIEE (10, 12). Moreover, HCN-channel dysfunction can also be acquired following experimental febrile seizures (13, 14) or in chronic epilepsy (15), affecting, for example, the epigenetic regulation through neurogenesis repressor element 1 (RE1)-silencing transcription factor (REST)/neuron-restrictive silencer factor (NRSF) (16).

While the contribution of HCN channels to biophysical properties of adult neurons is well established (17), their roles in brain development are mostly unknown. During postnatal rodent brain development, HCN channels undergo substantial and subunit-specific developmental regulation (7, 18–20). The four different pore-forming alpha subunits (HCN1 to HCN4) assemble into homo- or heteromeric tetramers (17). HCN channels are active at subthreshold membrane potentials and mediate *I*^h^, a voltage-dependent cation current activated by membrane hyperpolarization, which, in turn, depolarizes the membrane potential toward the action potential threshold and, thus, contributes to the dynamic control of the resting membrane potential and subthreshold integration (17). When active in stem cells, HCN channels could contribute to membrane depolarization or membrane voltage oscillations throughout the cell cycle. As pharmacological inhibition of HCN channels in cultured embryonic stem cells affected proliferation and differentiation (21, 22), neural stem and progenitor cells could likewise be affected by impaired function of HCN channels (23).

In this study, we sought to address whether the loss of HCN-channel function would affect brain development. Using transgenic mouse lines with altered HCN-channel activity in the embryonic telencephalon and pharmacological inhibition of *I*_*h*_ in cultured embryonic rat cortical neural stem cells (NSCs), we show that the loss of HCN-channel function impaired cell-cycle progression and proliferation and caused a pronounced microcephaly phenotype. We, thus, propose a mechanistic link between *HCN*-channel dysfunction and microcephaly in mice that may contribute to pathological cortical development in patients with *HCN* channelopathies.

## RESULTS

### Expression of HCN subunits during brain development

The four HCN subunits, HCN1-HCN4, are expressed at various levels in the mammalian central nervous system, and are known to form homomeric or heteromeric channels (17, 24). In the rat hippocampus, expression of the HCN1, 2, and 4 subunits changes during postnatal development. HCN1 and HCN2 are increasing, while HCN4 is decreasing with maturation (7, 18, 19). Therefore, we first determined the subunit-specific spatio-temporal expression patterns of HCN1 to HCN4 in the mammalian forebrain, focusing on embryonic development. Analysis of mRNA levels throughout pre- and postnatal mouse forebrain development revealed a monotonically increasing expression of *Hcn1* and *Hcn2*, whereas *Hcn3* and *Hcn4* expression levels peaked at embryonic day (E) 15, followed by a decrease (Fig. 1 *A*, *SI Appendix*, Fig. S1 *A-C*, and *A’-C’*). *Hcn3* was the most abundantly expressed subunit at E13-E18. *Hcn4* was the second highest expressed *Hcn* transcript at E13 that was present at levels comparable to that seen in *Hcn1* and *Hcn2* at E15-E18, after which it declined. Immunoblots of E15 mouse embryonic brain lysates detected HCN3 and HCN4 subunit proteins, while those of HCN1 and HCN2 were below the detection limits of the antibodies used (Fig. 1 *B*). Prenatal expression of the HCN subunits in developing human brain investigated by exon array hybridization of cDNAs generated from total RNA of frozen tissue samples (25) was detected during the earliest analyzed time period of late embryonic development (four to eight weeks post conception). As in developing mouse brain, HCN3 and HCN4 peaked during early to mid-fetal periods. Except for the thalamus, all brain regions showed a similar time course of expression (*SI Appendix*, Fig. S2). HCN1 and HCN2 increased at mid-gestation and reached almost adult-like levels around birth and during the first year of life (*SI Appendix*, Fig. S2).

**Figure 1.**
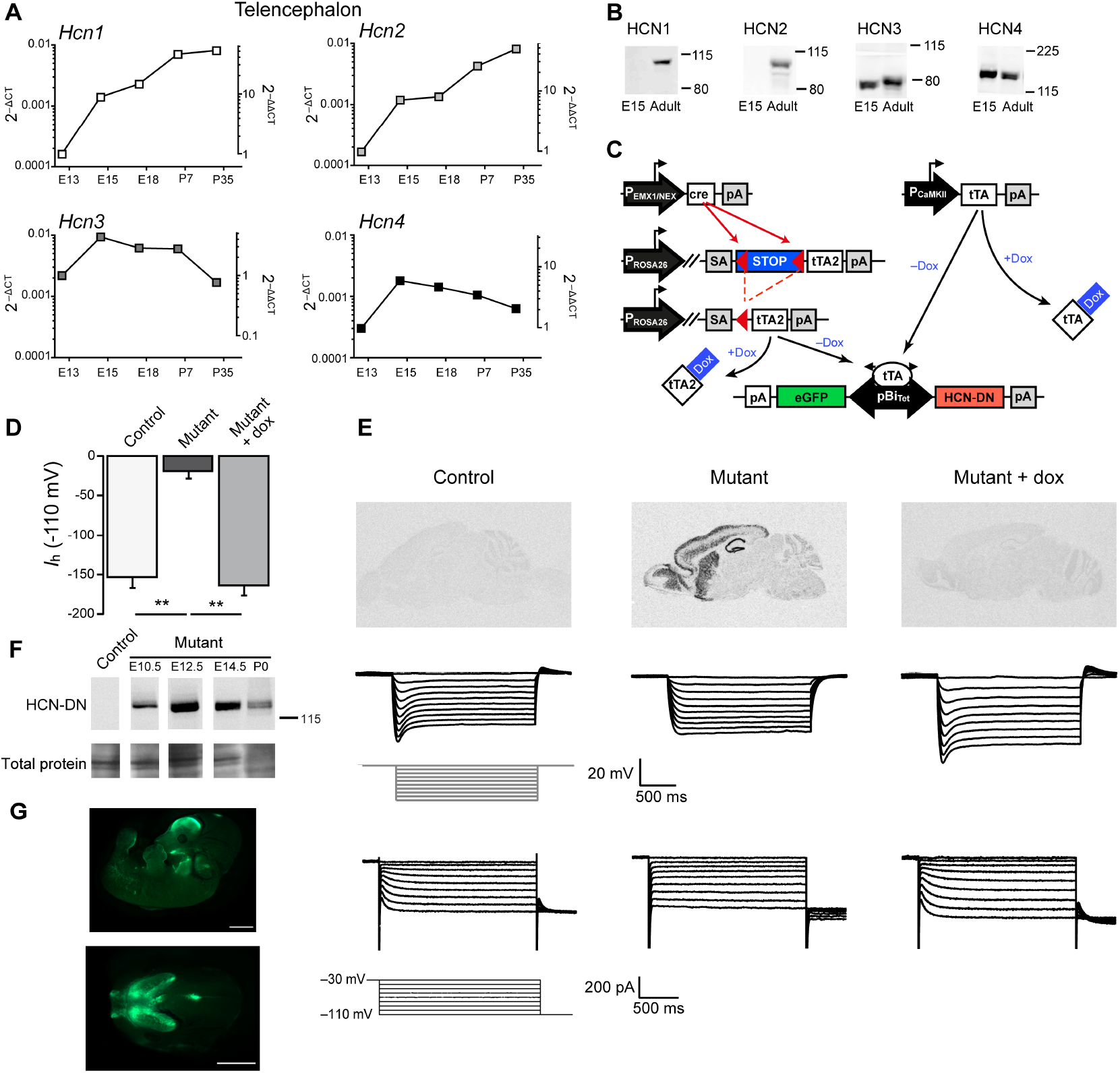
Embryonic HCN-subunit expression in mouse forebrain and dominant-negative transgenic strategy. **(A)** Quantitative RT-PCR analysis of *Hcn1-Hcn4* mRNA expression levels in mouse forebrain normalized to glyceraldehyde-3-phosphate dehydrogenase (GAPDH) mRNA (2^-ΔCT^, left y axis) and GAPDH and of the respective HCN subunit expression level at E13 (2^-ΔΔCT^, right y axis). **(B)** Immunoblot analysis of total brain lysates with HCN subtype-specific antibodies at embryonic day E15 and at the adult age. **(C)** Schematic overview of the mouse lines expressing the dominant-negative HCN4 subunit (HCN-DN) in embryonic or adult mouse brains. The embryonic expression was controlled by the EMX1 or NEX promoters driving Cre-recombinase expression, which removed the floxed stop cassette preceding the tetracycline transactivator (tTA2). Postnatal expression of HCN-DN was driven by the CaMKII promoter. In the absence of doxycycline, HCN-DN, which has an N-terminal hemagglutinin (HA) tag, and eGFP were expressed from the bidirectional tetracycline-responsive promoter pBi_Tet_. **(D)** Summary statistics from voltage-clamp experiments shown in (E) revealed markedly reduced *I*_*h*_ amplitudes in HCN-DN-expressing mutant cells (**, p < 0.01; one-way ANOVA with Tukey’s post-hoc test). **(E)** Doxycycline and genotype-dependent transgene expression in the brain was tested in *in situ* hybridization experiments using HCN-DN-specific probes (top row). Traces representative of current-clamp (middle row), or voltage-clamp recordings (bottom row), from CA1 pyramidal neurons in acute slice preparations from adult control (n = 14 cells; 4 mice), mutant (n = 11 cells; 4 mice), and doxycycline (dox)-treated mutant (mutant + dox, n = 9 cells; 2 mice) CaMKIIα-HCN-DN mice. The protocols are shown below the control traces. **(F)** Immunoblot detection of the HA-tag of the HCN-DN transgene in tissue lysates from mutant EMX1-HCN-DN brains at E10.5, 12.5, 14.5, at postnatal day (P) 0, and at E10.5 in a control brain. **(G)** eGFP expression in the whole embryo and head of E12.5 EMX1-HCN-DN mutants. Scale bars 1mm. Data are presented as mean ± s.e.m; Experiments in (A, B, F) represent individual examples. Images in (G) are representative of 5 animals.

Transcriptome analyses of select neural progenitor subpopulations and of young neurons isolated from E14.5 embryonic mouse and human fetal neocortex at 13 weeks post conception (26) indicated that *HCN3* was the main HCN subunit detected in both species (*SI Appendix*, Fig. S1 *D, E*).

At single-cell resolution, the transcriptome analysis of the developing brains from E9-E13 mouse embryos (data set III (27)) revealed the presence of *Hcn1, Hcn3*, and *Hcn4* transcripts in different classes of progenitors (*SI Appendix*, Fig. S1 *F, G*). Although at low levels, *Hcn* subunits were detected in the cell cluster expressing early stem cell/apical radial glia cell markers (Pax6, Sox2, Mki67) in the apical radial glia-specific marker (Eomes/Tbr2, Neurog2)-expressing cell cluster, and in the cell cluster expressing the neuronal marker Tubb3 (TuJ1) (*SI Appendix*, Fig. S1 *F-H*). In late progenitors/immature neurons (NPC) and in oligodendrocyte progenitor cells (OPC), *Hcn4* and *Hcn3* were the most prevalent subunits, while *Hcn1, Hcn3*, and *Hcn4* were most prominent in neural stem cells (NSCs) (*SI Appendix*, Fig. S1 *G*). Interestingly, Cajal-Retzius cells showed the highest expression of *Hcn* transcripts (*SI Appendix*, Fig. S1 *G*).

Single-cell RNA sequencing (scRNA-seq) of primary cultured embryonic E14.5 rat cortical neural stem cells (NSCs) using two different technical platforms revealed the presence of *Hcn1-4* in about 8% of NSCs analyzed using the well-based counts (WaferGen platform), and of *Hcn3* and *Hcn4* in 2.4% of the cells using the drop-seq (10x Genomics platform) expression results (*SI Appendix*, Fig. S3 *A, B*). In line with these findings, immunoreactivity against HCN3 was most frequently detected in rat NSCs (12.2±0.8% of cells), followed by HCN4 (9.2±1.3%) (*SI Appendix*, Fig. S3 *C*), which was also detected in cortical sections from E13 embryos (*SI Appendix*, Fig. S1 *I*), and HCN1 (6.2±2.2%) (*SI Appendix*, Fig. S3 *C*). Using the patch-clamp technique, we detected *I*_*h*_-like currents in only a small subset of cultured NSCs (*SI Appendix*, Fig. S3 *D*), which, together with the mRNA expression data, suggests the presence of a heterogeneous cell population, or only a short-lived presence of HCN channels in the cell membrane. Together, these results provide evidence for embryonic expression of HCN subunits and indicate a potential role for HCN1, HCN3, and HCN4 subunits in the developing telencephalon. As HCN subunits assemble into homo- or heteromeric tetramers (17), the presence of several HCN subtypes suggested that pharmacological or genetic dominant-negative strategies targeting HCN-channel function in a subunit-independent manner were most likely to unmask a role for *I*_*h*_ in embryonic brain development. In contrast to previous mouse models with single HCN-subunit knockouts (28–31), we generated conditional transgenic Tet-Off mouse lines expressing the hHCN4(p.Gly480Ser) subunit (HCN-DN) and an eGFP reporter in mouse forebrain (Fig. 1 *C*).

### Loss of HCN-channel function in forebrain NPCs causes microcephaly

Expression of HCN-DN was designed to ablate *I*_*h*_ by the formation of non-conducting heteromeric channels with the endogenous HCN1-4 subunits (*SI Appendix*, Fig. S4), which was prevented by the addition of doxycycline to the animals’ drinking water or chow (Fig. 1 *C, E*). This dominant-negative strategy was previously used by us to study HCN- or Kv7-channel function in the heart (32) or brain (33, 34), respectively. In adult mouse hippocampal CA1 pyramidal neurons (targeted by utilizing the CaMKII promoter) (Fig. 1 *C*), which predominantly express HCN1 and HCN2 subunits, the presence of HCN-DN led to a robust and doxycycline-dependent ablation of *I*_*h*_ (Fig. 1 *D, E*). Embryonic expression of HCN-DN was achieved under the control of the EMX1 promoter (EMX1-HCN-DN mutants), which started expression confined to the dorsal telencephalon at approximately E9.5 (Fig. 1*F*). On embryonc day 12.5 (E12.5), postnatal day 0 (P0), and P21, eGFP expression was mainly restricted to the telencephalon (Fig. 1 *G*, Fig 2 *A*), with the exception of a small region in the mesencepahlic vesicle and partial skin epression (Fig. 1*G*). The visible expression outside of the nervous system is in agreement with the known EMX1 expression (35). EMX1-HCN-DN mutants were viable but showed a decrease in body weight at birth (P0), which was even more pronounced at P21 (Fig. 2 *B, C*) and likely to be caused by developmental delay and reduced fitness as compared to their control littermates. At both time points, the forebrain morphology was severely altered, including a marked reduction in the cortex size and the absence of hippocampal pyramidal layers (Fig. 2 *D, E*). Using magnetic resonance volumetry, we quantified brain and ventricle volumes at P0 and P21. We found a marked reduction in brain volume at both ages and an increase in ventricle volume at P21 (Fig. 2 *G-J*). The restriction of HCN-DN expression to postnatal development by the application of doxycycline until P0 prevented the changes in body weight and the development of a microcephalic phenotype (Fig. 2), ruling out a postnatal or a positional effect of the transgene.

**Figure 2.**
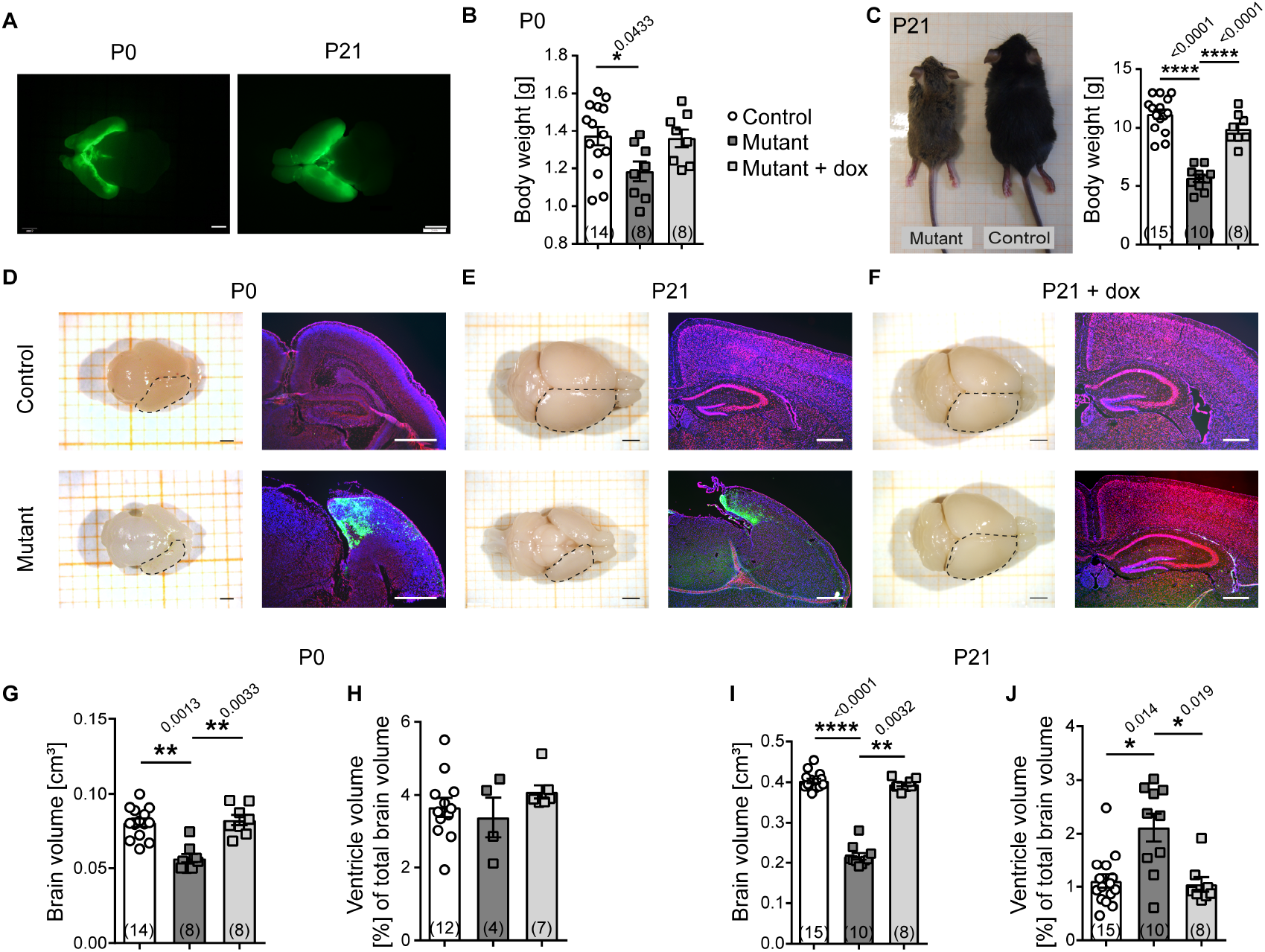
Development-dependent structural changes in *I*_*h*_-deficient mice. **(A)** eGFP expression in P0 and P21 EMX1-HCN-DN mouse brains is restricted to the forebrain. Body weights of **(B)** P0 and **(C)** P21 EMX1-HCN-DN mutant mice were lower than those of control animals and mutants on doxycycline until P0 (Mutant + dox) (one-way ANOVA with Tukey’s post-hoc test). Macroscopic top views of explanted brains and fluorescent Nissl staining (red) of coronal brain slices of EMX1-HCN-DN mice at **(D)** P0, **(E)** P21, and of **(F)** P21 mutant + dox. Slices were counterstained with DAPI (blue), eGFP is depicted in green. **(G)** Absolute brain volumes and **(H)** ventricle volumes as a fraction of the total brain volume of P0 EMX1-HCN-DN mice compared to those of control and mutant + dox animals. **(I)** Absolute brain volumes and **(J)** ventricle volumes as a fraction of total brain volume of P21 EMX1-HCN-DN compared to those of mutant + dox and control mice. (Kruskal-Wallis with post-hoc Dunn’s test.) Scale bars in P0 brains in (A) and (D) 1 mm, in P21 brains in (A, E, F) 2 mm, in sections 500 µm; Data is presented as mean ± s.e.m; n is given in parentheses; p values are above asterisks; Images in (D-F) right panel are representative of one experiment, in (A, D-F) left panel corresponding to n in (G–J).

### Early embryonic ablation of HCN or *I*_*h*_ inhibition in neural stem cells leads to decreased proliferation due to cell-cycle arrest in G1 phase

To analyze the mechanism underlying the marked microcephaly, we sought to investigate the critical processes of embryonic brain development that could impact brain size, such as neural progenitor cell proliferation, neuronal differentiation, apoptosis, and migration. At E12.5, most of the cells in the control or EMX1-HCN-DN cortex were Pax6-expressing radial glia cells (Fig. 3 *A*). As early as at this stage, a markedly reduced thickness of the mutant cortex was evident and accompanied by an increase in the apoptosis marker cleaved Caspase-3 and a decrease in the proliferation marker Ki67 (Fig. 3 *B-E*). The latter was not limited to embryonic stages but similarly found at P0, when the number of Ki67-immunoreactive cells in the ventricular and subventricular zones (VZ/SVZ) was reduced (Fig. 3 *F, G*). No increase was found in apoptotic cells, however. (*SI Appendix*, Fig. S5 *A, B*). A comparable number of proliferative cells was found in the sensorimotor cortex that likely reflected the ongoing gliogenesis in newborn mice (Fig. 3 *F-H*). To test whether the microcephaly phenotype was linked to the loss of HCN-channel function and not an artifact of the HCN-DN and eGFP transgenes, we used the same EMX1 promoter line as in EMX1-HCN-DN mice to co-express eGFP and a cAMP-insensitive HCN4-channel subunit (HCN4-573X), which, in contrast to HCN-DN subunits, does form functional channels (36). The resulting EMX1-HCN4-573X mice did not show structural changes (*SI Appendix*, Fig. S6) when analyzed at the age of 15 weeks.

**Figure 3.**
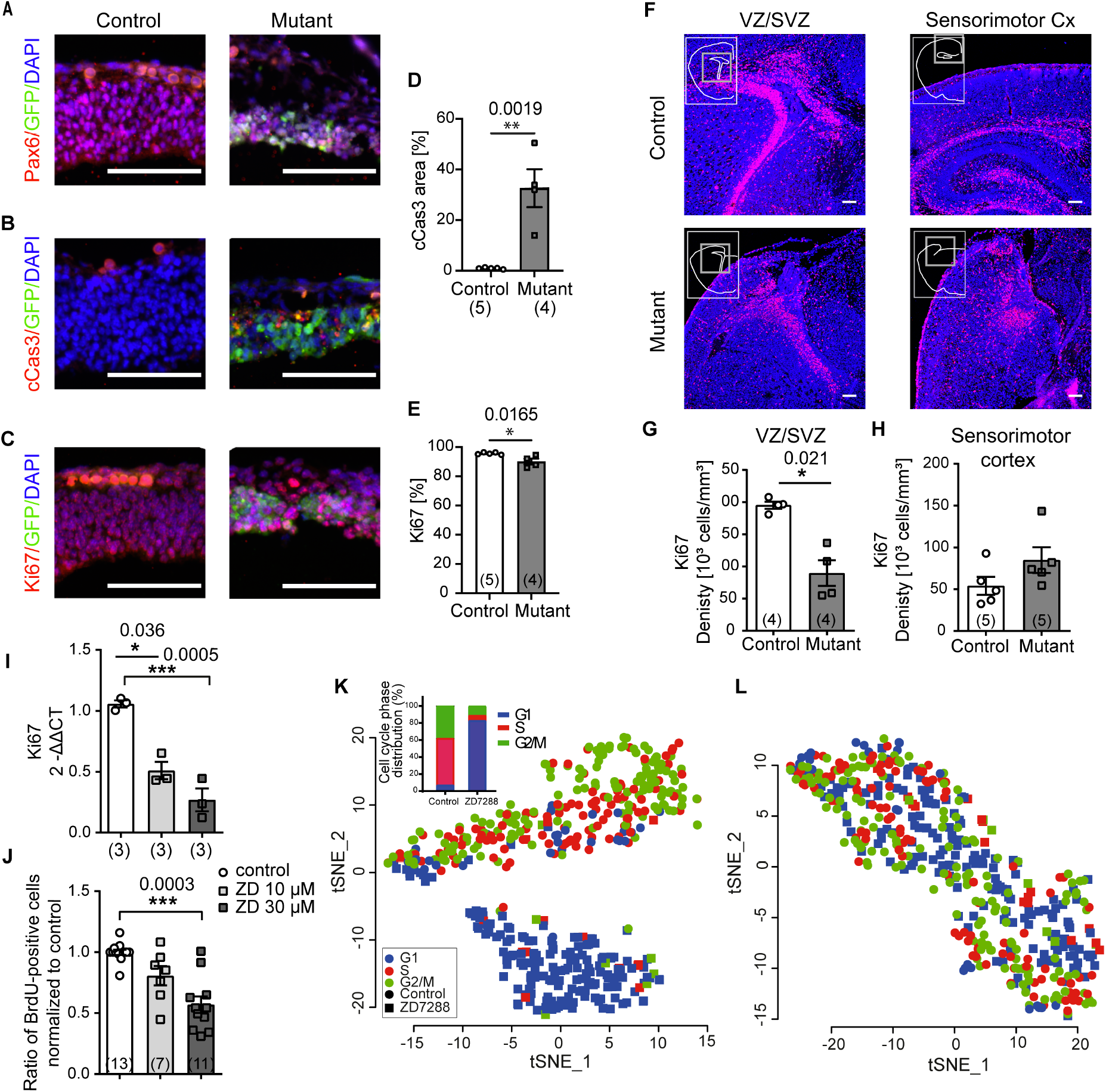
Impact of *I*_*h*_ blockade on proliferation and cell-cycle state. **(A-C)** Cortical sections of E12.5 control and EMX1-HCN-DN mutants stained for (A) radial glia cells (Pax6, red), (B) apoptotic (cleaved Caspase-3; cCas3, red), and (C) proliferating cells (Ki67, red), eGFP is depicted in green, DAPI in blue. **(D)** Quantification of cCas3-positive area in percentage of total analyzed area (unpaired t-test). **(E)** Quantification of Ki67-positive cells in percentage of DAPI-positive cells (unpaired t-test). **(F)** Images representative of Ki67 (red) staining of the VZ/SVZ and the sensorimotor cortex in P0 control mice and EMX1-HCN-DN mutants, DAPI is shown in blue. **(G, H)** Quantification of Ki67 expression (unpaired t-test). **(I, J)** Analysis of Ki67 mRNA level (I) and BrdU incorporation (J) in rat NSCs treated with ZD7288 (10 µM and 30 µM; Ki67: one–way ANOVA with Tukey’s post-hoc; BrdU: Kruskal-Wallis with post-hoc Dunn’s test). **(K)** Two-dimensional, tSNE projection of scRNA-seq data from vehicle-treated control (circles, 281 cells), and ZD7288-treated (30 µM, square, 190 cells) rat NSCs. The cell-cycle phase of each cell, which was determined from its gene expression profile, is indicated by the legend (lower left). The inset shows the relative distribution of cell-cycle phases in the vehicle-treated (control) and ZD7288 -treated NSC groups, which were significantly different from each other (p < 0.0001, Chi-square). **(L)** tSNE projection of the same NSCs as in (K) after regressing out the variance explainable by cell-cycle gene transcripts. Scale bars 100 µm; Data are presented as mean ± s.e.m; n is given in parentheses; p values are above asterisks; Images in (A, B, C) are representative of 5/4 animals, (F) are representative examples of (G, H)

We next generated mice in which HCN-DN expression was initiated in Nex (*Neurod6*)-expressing cells using the NEX-cre promoter line (NEX-HCN-DN mutants), which initiates cre-dependent gene expression starting at E10.5 in the SVZ, intermediate zone, and cortical plate (37, 38). Unexpectedly, the brain morphology of NEX-HCN-DN mice was equally unaffected (*SI Appendix*, Fig. S7) as in EMX1-HCN4-573X mice. This result prompted us to characterize *Emx1* and Nex/*Neurod2* expression in the mouse embryonic scRNA-seq dataset (27) in more detail. In contrast to *Emx1*, Nex/*Neurod6* expression was virtually absent from the NSC-cell cluster and, only to a minor extent, overlapped with the expression of *Mki67* (*SI Appendix*, Fig. S8 *F, G*). These expression data indicate predominant Nex/*Neurod6* expression in non-proliferating cells in the developing brains of E9-E13 embryos, which is in line with previous findings that cre-positive cells in NEX-cre mice did not incorporate bromodeoxyuridine (BrdU) at E16 (38). These results indicate that microcephaly in EMX1-HCN-DN mice is linked to the loss of HCN-channel function in NSCs of the VZ, thereby supporting the notion that functional HCN channels are needed for proper cell-cycle progression in NSCs.

To further test our hypothesis, we employed a different experimental system and analyzed the early developmental processes in NSCs from E13.5 rat cortices using the pharmacological *I*_*h*_ inhibitor ZD7288. Both the quantification of the mRNA levels of the proliferating cell marker Ki67 and percentage of BrdU-incorporating cells (cells in S phase) showed a ZD7288 dose-dependent decrease in proliferation (Fig. 3 *I, J*). Transcriptome analysis of scRNA-seq experiments of cultured NSCs displayed a clear separation between control and ZD7288-treated cell clusters (Fig. 3 *K*). To link transcriptomic data to biological pathways, we performed enrichment analysis of differentially regulated KEGG pathways revealing *DNA replication* and *cell cycle* as the two most affected pathways (Table 1). Determination of the cell-cycle phases showed that the majority of ZD7288-treated NSCs (82.8%) were in G1 as compared to control cells (7.0%), which were predominantly in the S and G2/M phases (Fig. 3 *K* inset). After regressing out the variance explainable by the cell-cycle marker genes from the dataset, the cells did no longer form clusters specific to their treatment (Fig. 3 *L*). The complementary, drop-seq-based NSC scRNA-seq data set II with higher cell numbers at the cost of losing visual single-cell shape control and having a lower sequencing depth, confirmed, although to a lower extent, G1 accumulation after pharmacological *I*_*h*_ inhibition (*SI Appendix*, Fig. S9). These results suggest the mechanism underlying the decrease in NSC proliferation is likely to be an *I*_*h*_ inhibition-induced increase in the length of the G1-cell-cycle phase that severely affects cell-cycle progression.

**Table 1.**
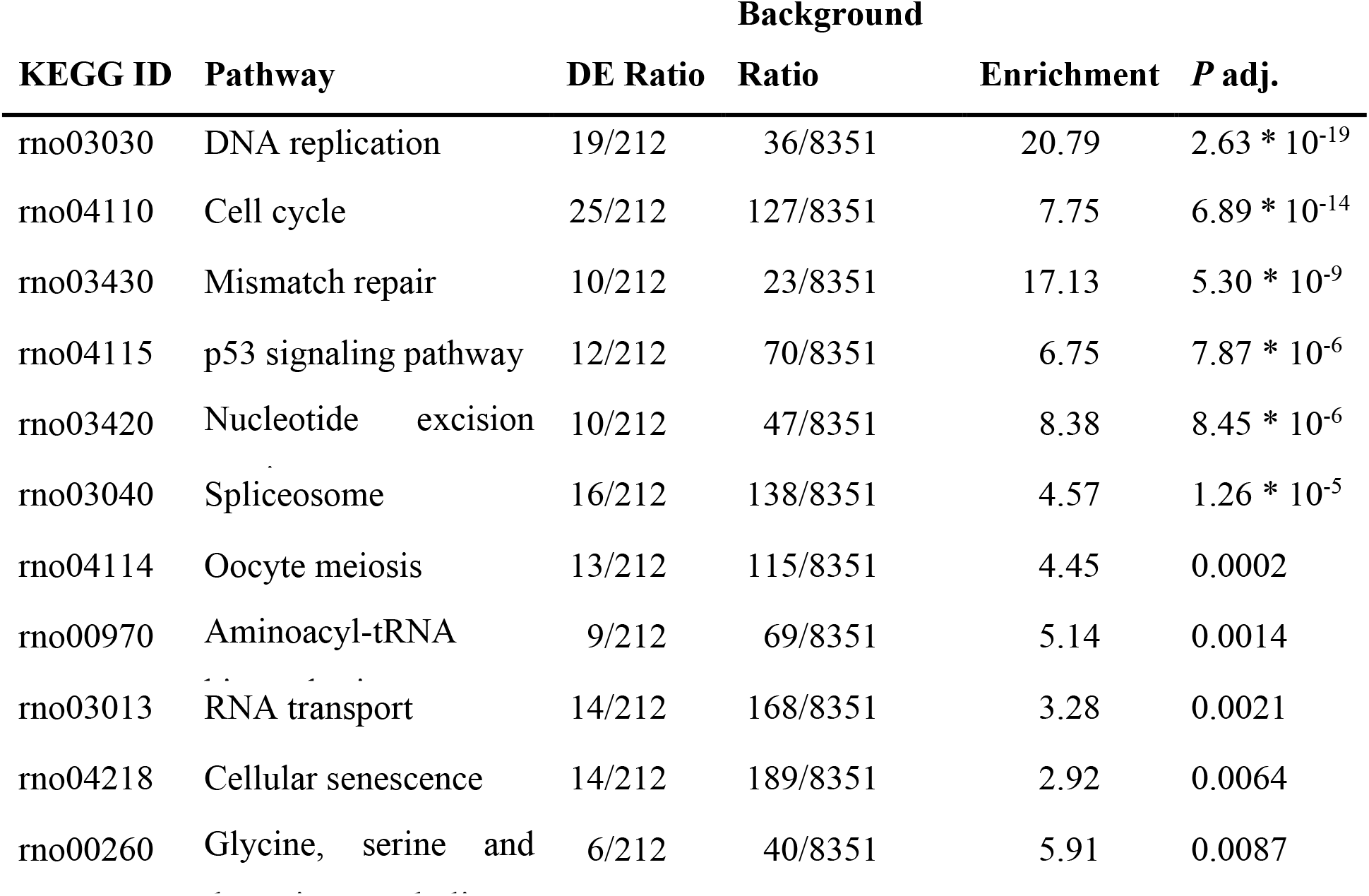
Gene set enrichment analysis of KEGG Pathways

In both datasets, HCN subunit-expressing untreated or ZD7288-treated NSCs did not cluster in clearly identifiable subclusters or cell types and, thus, did not contribute to the identification of a specific progenitor type (*SI Appendix*, Fig. S9 *B, E*). The increased cell numbers allowed cell cycle phase-specific differential gene expression analysis to be performed and revealed differential regulation of genes involved in cell cycle progression, associated with microcephaly, cortical neurogenesis, and stem cell differentiation (*Asns, Psat1*) (*SI Appendix*, Dataset S1) (39–41). Of note, we found a substantial overlap of differentially regulated genes after *I*_*h*_ inhibition in the rat NSC culture with an early developmental transcriptomic signature in cortical extracts from the Elp3 conditional knockout (Elp3cKO) microcephaly mouse model with impaired indirect neurogenesis due to endoplasmic reticulum (ER) stress and unfolded protein response (UPR) (42). In comparison, 13 out of 82 and 45 out of 405 differentially regulated genes in dataset I and dataset II, respectively, were also differentially regulated in the Elp3cKO mice (*SI Appendix*, Dataset S1).

### Effects of genetic or pharmacological *I*_*h*_ inhibition on differentiation, migration, and cell death

Next, we investigated the levels of the glial proteins GFAP and CNPase and the neuron-specific TuJ1 as markers of differentiation in brains from EMX1-HCN-DN mice and in cultured neural progenitors after ZD7288-mediated block of *I*_*h*_. In P0 mouse brains, we found higher immunoreactivity levels for GFAP (astrocytes) and CNPase (oligodendrocytes) in EMX1-HCN-DN mutants compared to those in control littermates, but no difference in TuJ1 levels (Fig. 4 *A-D*). In cultured primary rat NSCs, differentiation was initiated by FGF2 withdrawal, which induced a comparable decrease in the ratio of Sox2-positive cells in vehicle- or ZD7288-treated cells over time (Fig. 4 *E*). While the proportion of cells expressing the marker for both immature and mature neurons TuJ1 was decreased by ZD7288 five days after FGF2 withdrawal (Fig. 4 *F*), the ratios of the markers for astrocytes (GFAP) and oligodendrocytes (CNPase) were not significantly altered upon *I*_*h*_ inhibition (Fig. 4 *G, H*). Furthermore, neither HCN-DN expression in neural precursors induced at E15 by *in utero* electroporation nor ZD7288-mediated block of *I*_*h*_ in cultured NSCs affected cell migration (*SI Appendix*, Fig. S5 *E-G*). Unlike in sections from E12.5 embryonic cortex (Fig. 3 *B, D*), we did not find evidence for increased apoptosis/necrosis in the rat NSC cultures treated with ZD7288 (*SI Appendix*, Fig. S5 *C, D*). Likewise, Ca^2+^ handling of NSCs was not altered in a systematic way after application ZD7288 (*SI Appendix*, Fig. S5 *H, I*).

**Figure 4.**
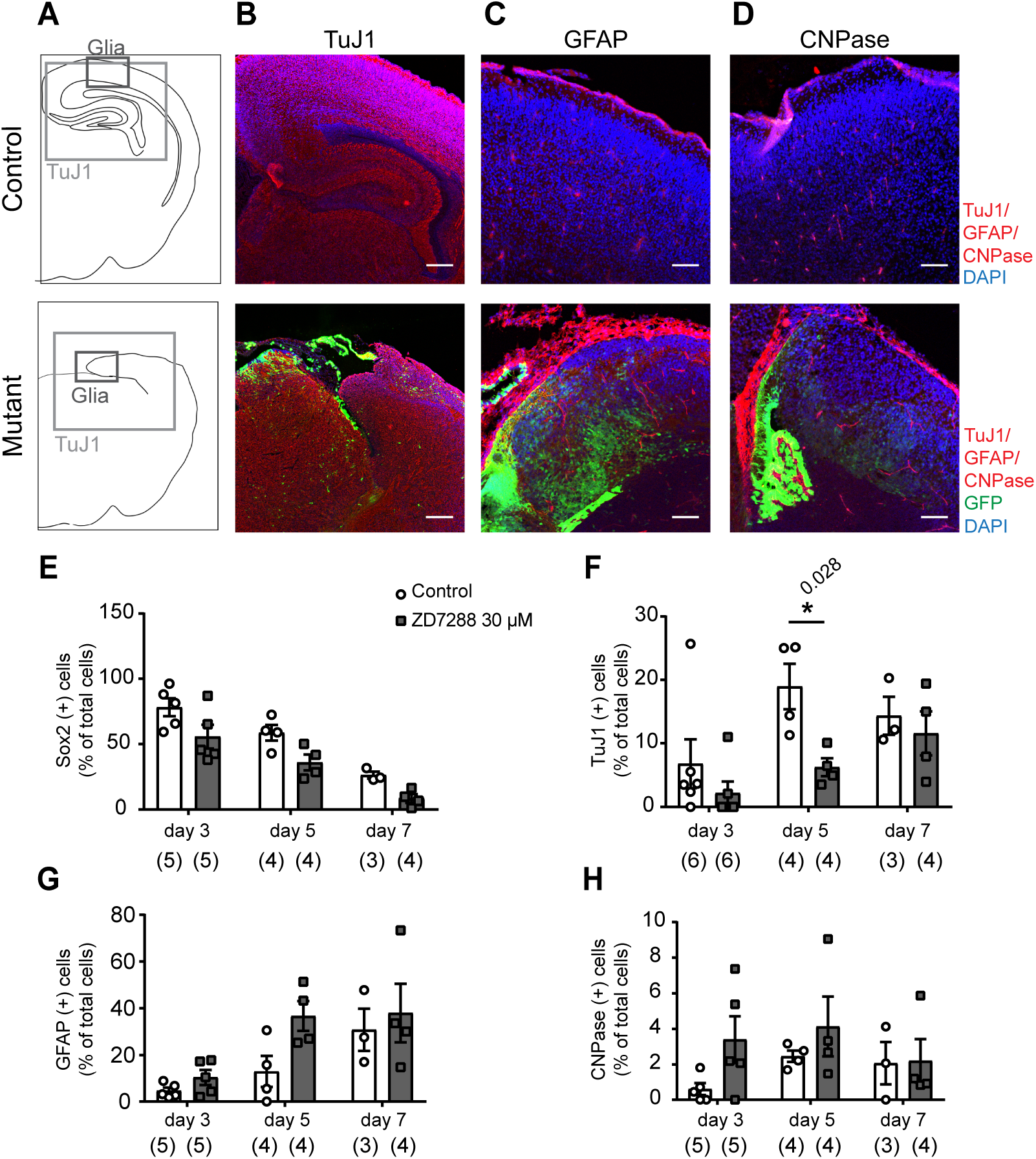
Influence of *I*_*h*_ ablation on differentiation in cultured NSCs and P0 mouse brains. **(A)** Schematic drawing of coronal sections with the region highlighted that is shown in the following sections. Immunofluorescence detection of **(B)** neurons (TuJ1), **(C)** astrocytes (GFAP), and **(D)** oligodendrocytes (CNPase) in the EMX1-HCND-DN (mutant) neonatal brain compared to those in controls. (E–H) Cortical stem cells differentiated in the absence of FGF2 were analyzed at days 3, 5, and 7 after FGF2 withdrawal (Mann-Whitney test). **(E)** The percentage of cells expressing the neural stem cell marker Sox2, **(F)** the neuronal marker Tuj1, **(G)** the astrocyte marker GFAP, and **(H)** the oligodendrocyte marker CNPase. For images representative of the differentiated cell cultures, see **SI Appendix, Fig. S10**. Scale bars (B) 200 µm, (C, D) 100 µm; Data are presented as mean ± s.e.m; n is given in parentheses; p values are above asterisks; Images in (B) are representative of one experiment and (C, D) of three similar experiments.

## DISCUSSION

Together, our findings establish a hitherto unknown role for HCN channels in the proliferation of neural stem and progenitor cells by affecting cell cycle progression and the exit from the G1-cell-cycle phase. The pharmacological approach, in combination with the genetic ablation of *I*_*h*_, suggests an ionic current-dependent mechanism underlying this phenotype. This notion is further supported by the analysis of the EMX1-HCN4-573X mouse line, which expressed a cAMP-insensitive, but otherwise functional HCN4 subunit in the embryonic telencephalon and did not show morphological alterations of the forebrain structure (*SI Appendix*, Fig. S6). Likewise, NEX-HCN-DN mutants, in which cre-dependent transgene expression starts only about one day later than in EMX1-cre mice in the SVZ, intermediate zone and cortical plate (37, 38) displayed normal brain morphology. Unlike *Emx1*, Nex/*Neurod6* expression was virtually absent from the NSC and proliferating cell clusters, indicating that Nex/*Neurod6* was predominantly expressed in non-proliferating cells in the developing brains of E9-E13 mouse embryos (*SI Appendix*, Fig. S8 *F, G*). The results obtained using these complementary genetic strategies provide evidence that the marked microcephaly was caused by the loss of HCN-channel function in NSCs of the VZ, rather than by off-target effects of the transgenic approach, thereby supporting the notion that functional HCN channels are needed for proper cell-cycle progression in NSCs.

Although we and others (23) did not find marked ZD7288-induced changes in the differentiation of cultured NSCs, in addition to the lower ratio of immature neurons five days after FGF2 withdrawal (Fig. 4 *E-H*), we observed early expression of the glial markers GFAP and CNPase in sections from P0 EMX1-HCN-DN mutants (Fig. 4 *A-D*), suggesting that in these animals the switch between neurogenesis and gliogenesis might have prematurely occurred. In contrast, GFAP and CNPase expression levels in adult EMX1-HCN4-573X and NEX-HCN-DN mutants were indistinguishable from those in controls (*SI Appendix*, Figs. S6, S7). These results indicate that, in addition to — and perhaps as a consequence of — a direct effect on NSC proliferation, a dysfunction in HCN subunits might also result in a dysregulated balance between neurogenesis and gliogenesis. A recent study found that both the application of ZD7288, or of the less specific *I*_h_ blocker Cs^+^, onto NSC cultures reduced the fraction of cells showing spontaneous Ca^2+^ oscillations (23). Using slightly different experimental conditions such as a shorter observation time, we did not find systematic changes in Ca^2+^ oscillations in our experiments with cultured rat NSCs upon ZD7288 application (*SI Appendix*, Fig. S5 *H-I*). ZD7288 is the most commonly used pharmacological *I*_*h*_ inhibitor that can show concentration-dependent off-target effects on T-type calcium channels or voltage-gated sodium channels (43–45). While voltage-gated calcium currents can be detected in neural stem and progenitor cells, only L-type calcium channels have hitherto been identified, which are not known to be blocked by ZD7288 (46–48). Furthermore, voltage-gated sodium channels are primarily associated with the differentiation, but not the proliferation, of neural progenitor cells (47–54). Therefore, both the dominant-negative transgenic strategy and the pharmacological block of *I*_h_ are likely to have affected the same HCN current-dependent mechanism.

HCN channels are part of an evolutionarily old and conserved family of ion channels (55) that are expressed in different cell types with proliferative potential (56, 57). A comparison of our HCN-subunit expression analyses with those of other studies and databases of embryonic mouse brain, stem cells, and induced human pluripotent stem cells has consistently revealed HCN3 to be the most abundant subunit (21, 23, 27, 56, 57). In addition to HCN3, the data also implicate HCN1 and HCN4 as potential HCN subtypes in the developing mouse and human fetal cerebral cortices (*SI Appendix*, Figs. S1, S2). The endogenous expression of all four HCN subunits and their potentially heteromeric channel assembly can be targeted by our dominant-negative approach in which *I*_*h*_ is inhibited in a subunit-independent manner. The low abundance of all HCN subunits and the marked phenotypes resulting from the loss of HCN-channel activity suggest a transient but essential function during the cell cycle. Prenatal expression of the HCN1 subunit, which is affected by mutations in the microcephalic EIEE patients (10, 11), started to increase at mid-gestation and was almost adult-like as early as at birth and during the first year of life ((25) and *SI Appendix*, Fig. S2). The expression pattern of *HCN1* in the developing human neocortex is, thus, in line with a role in late-embryonic/fetal and neonatal/early infantile brain development.

Brain malformation such as polymicrogyria is among the defects associated with ion channels, including NMDAR (here GRIN1, GRIN2B) (58, 59), or with voltage-gated sodium channels (SCN3A) (60). Previous publications show that the NMDA receptor is mainly expressed in migrating neurons, while the SCN3A gene is prominently expressed in the human-specific cell population of the outer radial glia cells and intermediate progenitor cells (60–62). This observation suggests an effect on mainly the migration of early neurons, or on specific proliferative populations, designed to expand the cell numbers for the advanced, gyrified human neocortex. In addition to causing polymicrogyria, different mutations in the NMDAR subunits can result in secondary microcephaly, which is potentially due to either apoptosis by non-migrated neurons or glutamate excitotoxicity caused by gain-of-function mutations (58, 59).

The differential gene expression analyses from our scRNA-Seq datasets comparing untreated and ZD7288-treated NSCs revealed a substantial overlap with genes identified in Elp3cKO mice, in which increased UPR in apical precursors affected indirect neurogenesis (via intermediate progenitors) (6). It is tempting to speculate that the marked apoptosis at E12.5 in our EMX1-HCN-DN mouse model is a downstream consequence of a slowed cell-cycle progression and subsequently increased UPR. The more pronounced microcephaly in our model as compared to the Elp3cKO model may indicate that the loss of HCN-channel function affected both direct and indirect neurogenesis by inducing, for example, also earlier progenitor depletion by apoptosis. Furthermore, the reduced proliferation of neural progenitors (Fig. 3) is attributable to a severe reduction in cell-cycle speed, which scRNAseq indicates is primarily due to G1 phase lengthening, ultimately causing depletion of the neural progenitor pool and, as a consequence, microcephaly. Loss of functional HCN channels in NSCs affected the membrane potential (23), which is more depolarized in cycling cells than in cell cycle-exiting cells (3). Together, the data indicate that *I*_*h*_, which contributes to rhythmic activity in many cell types likely including NSCs (23), also helps control the entry into or exit from the cell cycle in neural progenitors. This hypothesis is in line with recent findings showing that experimental hyperpolarization of the progenitor cell pool causes their premature shift to later developmental stages (6), thus highlighting the importance of bioelectrical membrane properties as determinants of cell proliferation and differentiation. While target genes of the Wnt signaling pathway were enriched among (mostly up-) regulated genes in the respective progenitors in that study (6), we did not find Wnt signaling target gene enrichment among differentially regulated genes in our NSC cultures after pharmacological *I*_*h*_ blockade, which argues for Wnt-independent mechanisms. As *I*_*h*_ inhibition with ZD7288 led to a reduction in proliferation not only in NSCs (our study and (23)) but also in embryonic stem cells (21, 22), HCN subunit-mediated currents may be part of a conserved mechanism that is not restricted to brain development. In this context, it is noteworthy that HCN1-channel expression is also controlled by REST/NRSF, the critical epigenetic regulator of neurogenesis. REST/NRSF is expressed in embryonic and neural stem cells and is essential for maintaining NPC fate by inhibiting neuronal specific genes (63, 64). Dysregulation of REST/NRSF through deficiency of its upstream regulator ZNF335 resulted in microcephaly in both humans and mice (65). The forebrain-specific, EMX1-cre-mediated knockdown of ZNF335 in mice disrupted progenitor cell proliferation, cell fate, and neuronal differentiation and — similar to the phenotype of EMX1-HCN-DN mice — caused a severe reduction in forebrain structure (65). Although REST/NRSF controls many genes (66), its dysregulation may result in a suppression of HCN1-channel expression at a critical cell cycle checkpoint and, thus, contribute to the observed phenotypes in mice and humans.

Although causality is difficult to prove in patients carrying *HCN1* mutations that are associated with EE and microcephaly (10, 11), the combined clinical and experimental data indicate that HCN-channel dysfunction-mediated impairment of neural progenitor proliferation is a mechanism that warrants further investigation into idiopathic brain malformations.

## MATERIALS AND METHODS

### Animals and husbandry

All experimental procedures were approved by the Behörde für Gesundheit und Verbraucherschutz of the City-State of Hamburg, and by the Landesamt für Natur, Umwelt und Verbraucherschutz Nordrhein-Westfalen, Germany. Animals were kept in Type II-long (Tecniplast) plastic cages under standard housing conditions (21 ± 2°C, 50-70% relative humidity, food, and water *ad libitum*, and nesting material provided). Mice were kept on an inverted 12:12 light:dark cycle.

In the EMX1-HCN-DN mouse lines mainly EMX1-Cre or HCN-DN negative animals were used as controls.

### Cloning of HCN constructs

Mouse HCN1 (mHCN1) and human HCN4 (hHCN4) cDNAs (GenBank Acc. NM_010408 and NM_005477) in the pcDNA1 vector were kindly provided by Benjamin Kaupp (Forschungszentrum caesar, Bonn, Germany). A hemagglutinin (HA) epitope tag (YPYDVPDYA) was attached to the N-terminus of the open reading frame of hHCN4, as described (67). The selectivity filter GYG motif was mutated to SYG by overlap PCR-based, site-directed mutagenesis resulting in hHCN4 (p.Gly480Ser) (HCN-DN). For two-electrode voltage-clamp experiments, mHCN1 and HCN-DN were cloned into the RNA expression vector pGem-HeJuel (68). For the generation of transgenic mice, HCN-DN was inserted in pBI (GenBank Acc. U89932, Clontech) containing eGFP. All coding sequences were verified by direct sequencing.

### Generation of Rosa26(loxP-stop-loxP-tTA2)^Isb^ mice

A floxed stop cassette (69) was inserted 5’ of the tTA2 coding sequence (Clontech) into the plasmid pcDNA3 (Invitrogen), and the resulting loxP-Stop-loxP-tTA2-BGH-poly(A) fragment was excised and inserted into the pBigT vector (70). Subsequently, the AscI–PacI insert of pBigT was isolated and inserted into the pROSA26-PA vector (70), resulting in the target vector pROSA26-stop-tTA2. The linearized targeting vector was electroporated into R1 ES cells, which were subjected to selection by geneticin (G418, Invitrogen). Homologous recombination was verified by Southern blot analysis of digested genomic ES-cell DNA.

### Generation of mice expressing HCN-DN and HCN4-573X

Transgenic mice carrying the pBI-HCN-DN/eGFP construct, Tg(Bi-tetO-HCN^p.G480S^, eGFP)^C Isb^), or mice carrying the pBI-HCN4-573X/eGFP construct, Tg(Bi-tetO-HCN^p.573X^,eGFP)^C Isb^) were generated by pronuclear injection using standard techniques. Founder mice and the resulting offspring were genotyped by PCR using ear or tail biopsies, backcrossed to the C57BL/6J (B6) background for more than ten generations, and crossed with promoter mice B6-Tg(Emx1^tm1(cre)Ito^) (35), B6-Tg(Nex-cre (NeuroD6^tm1(cre)Kan^)) (37), or B6-Tg(CaMKIIα-tTA)^1Mmay/J^ (71).

Transgenic mice of the B6-Tg(Emx1^tm1(cre)Ito^, Rosa26^tm1(loxP-stop-loxP-tTA2)Isb^, (Bi-tetO-HCN4^p.G480S^,-eGFP)^CIsb^) or B6-Tg(Emx1^tm1(cre)Ito^, Rosa26^tm1(loxP-stop-loxP-tTA2)Isb^, (Bi-tetO-HCN4^p.573X^,-eGFP)^CIsb^) mouse line were genotyped by PCR using the following corresponding primer sequences Cre: 5’ - AAA CGT TGA TGC CGG TGA ACG TGC - 3’, 5’ - TAA CAT TCT CCC ACC GTC AGT ACG - 3’ (214 bp); tTA: 5’ - CCA TGT CTA GAC TGG ACA AGA - 3’, 5’ - CTC CAG GCC ACA TAT GAT TAG - 3’ (597 bp) and for Hcn4: 5 ‘-GGC ATG TCC GAC GTC TGG CTC AC - 3 ‘, 5’ - TCA CGA AGT TGG GGT CCG CAT TGG - 3’ (350 bp). NEX-Cre transgenic mice were kindly provided by Prof. K.-A. Nave (Max Planck Institute for Experimental Medicine, Göttingen), genotyped as described (38), and crossed with the B6-Tg(ROSA26^tm(loxP-stop-loxP-tTA2)Isb^, (Bi-tetO-HCN^p.G480S^, eGFP)^C Isb^) mouse line. The CamK2α-tTA mouse line was genotyped by PCR using the following corresponding primer sequences tTA: 5’-CGC TGT GGG GCA TTT TAC TTT AG-3’ and 5’-CAT GTC CAG ATC GAA ATC GTC-3’ (450 bp).

Control animals lacked at least one (40% were negative for the EMX1-Cre allele, 49% were negative for HCN-DN) or two alleles (3% were negative for the tTA2 and HCN-DN alleles, 6% were negative for EMX1-Cre and HCN-DN). To suppress HCN-DN expression, mice received 200 mg doxycycline per kg chow or drinking water. For/In the EMX-HCN-DN +dox cohort, doxycycline was administered until the day of birth.

### Tissue preparation for histology

Mice were deeply anesthetized with a combination of ketamine and xylazine (100 mg/kg ketamine, Ketanest; Pfizer and 20 mg/kg xylazine, Rompun; Bayer) and transcardially perfused with phosphate-buffered saline (PBS), followed by phosphate-buffered 4% formaldehyde (PFA) solution (Histofix, Carl Roth). Brains were dissected and postfixed in 4% FA for 24 h, and cryoprotected in 15% and 30% sucrose in PBS overnight, respectively. Tissue was rapidly frozen using optimal cutting temperature compound (OCT) and cut into 14-µm-thick slices on a cryostat (CM 3050S, Leica). Tissue and slices were stored at -80°C.

Embryos were dissected and staged according to Theiler’s staging criteria at approximately embryonic day E12.5 (TS19-20). The embryonic heads were immersion fixed in 4% PFA solution and cryoprotected in 30% sucrose. The tissue was frozen in OCT and cut into 12-µm-thick slices.

### Fluorescent Nissl stain

The NeuroTrace® 530/615 Red Fluorescent Nissl Stain (Molecular Probes) was diluted 1:25 in PBS and applied according to the manufacturer’s protocol.

### Immunohistochemistry

Slides were washed three times in PBS for 5 min. Antigen retrieval was performed in 10 mM sodium citrate buffer (pH 9) at 70°C for 30 min for 2’,3’-cyclic-nucleotide 3’-phosphodiesterase (CNPase) and cleaved Caspase 3 staining. After cooling to room temperature in the buffer, slides were washed four times each in PBS for 5 min. Blocking of non-specific binding was performed at room temperature for one hour. For glial fibrillary acidic protein (GFAP) staining, the blocking solution contained 10% normal horse serum, 0.2% bovine serum albumin, and 1% Triton X-100 solution in PBS. For CNPase, cleaved caspase-3, Ki67, TuJ1, and GFP stains, the blocking solution contained 5% normal goat serum and 0.2% Triton X-100 in PBS. The primary antibody (Ki67: rabbit, AB15580, Abcam, 1:250; TuJ1: mouse, MAB1195, R and D Systems, 1:100; GFAP: mouse, clone GA5, MAB3402, Millipore, 1:500; CNPase: mouse, Clone 11-5B, C5922’ Sigma-Aldrich, 1:1000; cleaved Caspase 3: rabbit, AF835, R and D Systems, 1:2000; GFP: conjugated with A488, rabbit, A-21311, life technologies, 1:200) was diluted in PBS and applied overnight at 4°C. Brain slices were washed four times in PBS for 5 min and incubated for 2 hours in the secondary antibody (1:200 Alexa Fluor® 546/488 goat anti-mouse (A-11030), or in goat anti-rabbit (A-11035) IgG, Molecular Probes) in PBS at room temperature. HCN4 detection was performed using a phosphate buffer (PB) (12 M Na2HPO4, 5 M NaH2PO4, pH 7,4) for washing steps and the 2h-long block in 5 % Chemiblocker (Millipore), 0.5 % Triton X-100 in PB. The primary antibody was diluted 1:100 in PBS (HCN4: mouse, 75-150, NeuroMab). Further processing was performed as described above. Slides were washed four times for 5 min in PBS and embedded in DAPI Fluoromount-G (Southern Biotech).

Sections from E12.5 embryonic brains were heated in Tris-EDTA (5mM Tris-HCl, 1mM EDTA, pH 8) antigen retrieval buffer at 99°C for 20 min followed by a 20-min cool down at RT. After a 5-min PBS rinse, the sections were blocked in 10% normal donkey serum and 0.3% Tween20 solution in PBS for 1h. Primary antibodies (Pax6: rabbit, 901201, Biolegend, 1:1000; cleaved Caspase 3: rabbit, D175 (A51E), Cell Signaling, 1:2000; Ki67: mouse, 556003, BD Bioscience, 1:1000; GFP: goat, 600-101-215, Rockland, 1:2500) were accordingly diluted in 0.3% Tween20 in PBS and incubated overnight at 4°C. Secondary antibodies (Donkey anti-rabbit IgG (H+L) Alexa Fluor Plus 555, (A32794), Donkey anti-mouse IgG (H+L) Alexa Fluor Plus 555 (A32773), Donkey anti-goat IgG (H+L) Alexa Fluor Plus 488 (A32814), Invitrogen) were diluted 1:1000 in 10% normal donkey serum and 0.3% Tween20 in PBS and incubated for 2h at RT. For DAPI staining (D1306, Invitrogen), the slides were washed for 10 min in 1:50000 dilution in PBS and mounted with Prolong Glass Antifade Mountant (P36980, Invitrogen).

### Stereology

Stereological analyses were performed with a Nikon, or a Zeiss AX10, microscope equipped with StereoInvestigator software (Microbrightfield). For proliferation and apoptosis studies, four sections, spaced 84 µm, approximately between +2.31 mm and +3.75 mm from the olfactory bulbs, were analyzed and, according to the developmental mouse brain atlas, represent the sensorimotor cortex and the underlying VZ/SVZ (72). The optical dissector principle was used to estimate cell densities as described (73). The parameters of analysis were as follows: Guard space depth 2 μm, base and height of the dissector, 3600 μm^2^ and 10 μm, respectively, and the distance between the optical dissectors was 60 μm, objective 40× Plan-Neofluar^®^ 40×/0.75 (Zeiss). For the analysis of migration, 5 sections, 56 µm apart from each other, were studied. The electroporated area was subdivided into layers 2/3 (target region), layers 4-6, subplate and intermediate zone (migration region), and VZ/SVZ as a proliferative (starting) region. All electroporated cells in the regions of interest were counted.

### Image analysis E12.5 embryos

All images were taken using a Leica DM5500 B with a Leica DFC 360 FX camera. For analysis, FIJI Is Just ImageJ software (v1.53g) was used. All sections were matched on an anterior-posterior and on a medial-lateral axis. For Ki67 cell numbers DAPI and Ki67-positive cells were manually counted in a 140-µm-long fragment of the cortex. To establish the cleaved Caspase 3-positive area, a 140-µm-long area was drawn along the cortex and processed by using the ‘Triangle’ threshold settings and ‘Analyze Particle’ function (size: 0 – infinity, circularity: 0 – 1). The percentage of thresholded area to total region of interest was calculated. Four sections per embryo were analyzed.

### Magnetic resonance volumetry

For magnetic resonance imaging (MRI), neonatal animals were deeply anesthetized using a combination of ketamine (100 mg/ml) and xylazine (20 mg/ml). Subsequently, they were perfused with 7 ml PBS followed by 4% FA. Both solutions contained 3% gadolinium as an MRI contrast agent. The head was fixed overnight in FA gadolinium solution and stored in 3% gadolinium in PBS until measurement. The MRI was performed on a 7.0 T small animal scanner with a 3D protocol (ClinScan, Bruker). Neonatal animals were analyzed with turbo spin-echo [repetition time (TR) 200 ms, echo time (TE) 8.5 s, turbo factor 8, matrix 144 × 192 pixels, field of view (FOV) 11 × 14 mm, slice thickness 80 µm]. Three-week-old animals were measured alive under 1-2% isoflurane anesthesia (500 ml/min). Both the breathing rate and body temperature of the head-fixed animals were continuously monitored. The juvenile animals were imaged using a CISS sequence (TR 8.14 ms, TE 4.07 s, matrix 192 × 192 pixels, FOV 16 × 16 mm, slice thickness 90 µm). All further analyses were performed with Osirix (Pixmeo SARL). For quantitative analysis, the brain volumes included the midbrain, but not the cerebellum, because the respective boundaries were reproducibly identified both in the control and EMX1-HCN-DN mice.

### Immunoblots

Tissue samples were homogenized in a solution containing 330 mM sucrose, 20 mM 3-(N-morpholino) propane sulfonic acid (MOPS), 1 mM ethylenediaminetetraacetic acid (EDTA), and a proteinase inhibitor cocktail (Sigma-Aldrich). Membrane proteins were separated from cytosolic proteins by ultracentrifugation or a ProteoExtractR Transmembrane Protein Extraction Kit (Novagen). Protein concentration was measured with a bicinchoninic acid (BCA) protein assay (Thermo Scientific), according to the user manual. Proteins were separated using 12%-Bis-Tris sodium dodecyl sulfate-polyacrylamide gel electrophoresis (SDS-PAGE) in MOPS buffer. The blotting of proteins on the nitrocellulose membrane was performed in NuPAGE (Life Technologies) transfer buffer containing methanol. The total protein content on the membrane was determined using SERVAPurple staining (Serva) according to the manufacturer’s protocol. For immunostaining, all solutions were prepared with tris-buffered saline (TBS) containing 0.01% Tween20. The membrane was washed in TBS solution and blocked in 5% milk powder. The primary antibody was diluted 1:2000 for HCN-subunit detection and 1:500 for HA staining (HCN1: mouse, N70/28, NeuroMab; HCN2: mouse, 75-111, NeuroMab; HCN3: mouse, 75-175, NeuroMab; HCN4: mouse, 75-150, NeuroMab; HA: mouse, 12CA5, Roche) in 1% milk powder and incubated overnight at 4°C. The membranes were washed for 4 × 5 min and incubated with horseradish peroxidase (HRP)-coupled secondary antibody (1:500) for one hour at room temperature. For detection, Luminata(tm) Crescendo Western HRP Substrate (Millipore) was used.

### *In situ* hybridization (ISH)

To detect transgenic hHCN4-DN, a construct (polyA)-specific, α-32S-UTP (∼800 Ci/mmol)-labeled probe of 445-bp length within exon 8 of the hHCN4 (3148 – 3592 bp) was used. The ISH experiments were performed using standard techniques.

### Quantitative PCR (qPCR)

To analyze the HCN-subunit expression during different developmental changes, the TissueScan Mouse Developmental Tissue qPCR Array (Origen) was used. A TaqMan-PCR was performed with a commercially available primer and probe mix containing VIC™ labeled GAPDH probe (GAPDH: Mm99999915_g1), and FAM™ labeled HCN subunit probe (HCN1: Mm00468832_m1; HCN2: Mm00468538_m; HCN3: Mm01212852_m; HCN4: Mm01176086_m1; Applied Biosystems). A 7900HT Fast Real-Time PCR System (Applied Biosystems) instrument was used to perform the PCR. Data were analyzed with SDS 2.3, RQ Manager 1.2, and DATA Assist v2.0 (Applied Biosystems). Individual reactions were normalized to co-amplified GAPDH. For illustration, all time points were normalized to E13 of the respective HCN subtype.

### RNA-seq

The detailed methods of the cell type-specific transcriptome analysis were published elsewhere (26), and the data set was reanalyzed for HCN-subtype expression levels. To identify the HCN subtypes, the sequences were aligned with the mouse reference genome (mm10) and the human genome (hg19). The data are shown in fragments per kilobase per million reads (FPKM) with a threshold of 1.

Single-cell RNA-seq analysis of cultured NSCs isolated from the cortex of E14.5 Sprague-Dawley rats (NSC001, R&D systems; cultured according to the manufactures’ protocol) was performed according to the manufacturers’ protocols with two independent methods using either a well-based approach with four biological and one technical replicate using the WaferGen ICELL8 single-cell system (data set I) or a drop-seq approach with the 10x-chromium system (data set II, 10x Genomics). For the ICell8 approach, the indexed amplified cDNA was collected using a collection kit (WaferGen) and centrifuged at 3220 g for 10 minutes at 4°C. After concentrating the product with a DNA Clean & Concentrator Kit (Zymo Research), the cDNA was purified with 0.6X AMpure XP beads (Beckman). The quality of the cDNA was assessed on a Tape 12 Station (Agilent). A total of 1 ng purified cDNA was used as starting material for the library preparation. The cDNA was tagmented by Nextera XT transposome, and adapter sequences were successively added to the end of the fragments. In a subsequent step, amplification was carried out by PCR. The library was purified using a 1:1 ratio of AMpure XP beads, followed by dual SPRI (Solid Phase Reversible Immobilization) treatment.

The libraries from the 10X experiment were prepared with the Chromium Single Cell 3’ Library & Gel Bead kit v2 and i7 Multiplex kit (10X Genomics).

Finally, the quality and molar concentration of the library were assessed on a Tape Station. Sequencing was performed on an Illumina HiSeq4000.

For data set II, Cell Ranger Suite (v3.0.2, 10X Genomics) was used to perform sample multiplexing, barcode processing, and generation of a single-cell gene UMI (unique molecular index) count matrix.

### Single-cell transcriptome analysis

In total, the read count matrix of dataset I contains 129,136,294 reads, each read with one out of 510 molecular barcodes. After quality control of the reads using FastQ Screen (http://www.bioinformatics.babraham.ac.uk/projects/fastq_screen/), we obtained distinct RNA profiles for 510 barcodes, including five positive controls (total RNA from bulk samples of non-treated cells) and five negative controls (empty-well samples). After removing the controls, we obtained a matrix of 500 barcodes (corresponding to cells) times 23,425 ENSEMBL IDs (corresponding to genes).

All barcodes were required to comprise at least 200 different detectable genes (at least one count). This requirement was met for all barcodes. Subsequently, we removed the genes that were detected (at least one count) in fewer than seven cells. This left us with 14,588 genes. Based on the distribution of total counts per barcode, we removed all cells with more than 2^20^ counts because they potentially originated from wells containing multiple cells (*SI Appendix*, Fig. S11).

Dataset II contained 849,391,962 total reads from 4,691 cells, with a total of 25,399 detected genes. For further analysis, we considered only cells expressing 2,000 – 6,000 genes (with at least one UMI count) and genes expressed in at least three cells. We additionally removed cells containing more than 11% mitochondrial gene counts. The final UMI count matrix consisted of 3,492 cells and 13,802 genes.

All downstream analyses were performed using in-built functions from the Seurat Bioconductor package (74). Seurat assigns a cell-cycle phase (G1, S, G2/M) to each cell. To this end, we had to provide a list of cell-cycle marker genes for S and G2/M (*SI Appendix*, Dataset S2) and performed t-SNE mapping to a 2-dimensional space before and after cell cycle gene removal using the RegressOut function in Seurat. Differential gene expression analysis between groups was done using the Wilcoxon rank-sum test. Enrichment analysis of differentially regulated KEGG pathways was done using ClusterProfiler (75), and we employed TopGO (76) for Gene Ontology enrichment analysis. Dataset III was downloaded from http://mousebrain.org/downloads.html (27) and contains all scRNA-seq data from forebrain cells prepared from E9.0 – E14.0 embryos (Forebrain: E9.0, E10.0, E11.0; Forebrain dorsal: E12.0, E12.5; Forebrain ventral: E12.0, E13.0). The original dataset contained scRNA-seq data from 292,495 total cells with total 31,053 genes detected. After selection with respect to brain region and age, subsequent quality control removed or cells with less than 200 or more than 6,000 genes per cell detected. Cells with a mitochondrial transcript content of >15% were also removed. The final library contained 47,263 cells with a total of 31,053 genes detected with a 6,153 median UMI counts per cell. The cell-type assignment was based on molecular marker expression (*SI Appendix*, Table S1) to detect the major cell types present in late embryonic mouse brain. For the counting of *Hcn1-4* isoform-expressing cells, at least one UMI count per cell had to be detected.

### Cell culture

For primary neural stem cell (NSC) monolayer culture, rat cortices were derived from embryonic day 13.5, as described previously (77). Human recombinant fibroblast growth factor (human FGF2) (10 ng/ml, Invitrogen) was added to Dulbecco’s Modified Eagle’s Medium (DMEM)/F-12 (Gibco), supplemented with L-glutamine, N2-supplement, penicillin/streptomycin, and sodium pyruvate (Gibco). After two passages, 2 × 10^4^ cells were re-plated in chamber slides as monolayers in the presence of FGF2 for further experimental procedures. To assess the proliferation potential, migratory behavior, and cytotoxic effects, NSCs were treated with the *I*_*h*_ blocker ZD7288 (Sigma-Aldrich) at concentrations of 10 µM and 30 µM. RNA extraction experiments or assays were performed after 18 hours of incubation. Differentiation was initiated by the withdrawal of FGF2, and ZD7288 (30 µM) was added every 24 hours. After 3, 5, and 7 days following FGF2 withdrawal, NSCs were fixed and immunocytochemically stained.

### *In vitro* proliferation assays

For proliferation analysis, qPCR for Ki67 was performed. The GeneUp Total RNA Mini Kit (Biotechrabbit) was used for extracting the RNA from cortical stem cells. RNA amount was measured with the GeneQuant 1300 and transcribed into cDNA using the QuantiTect Reverse Transcription Kit (Qiagen). qPCR was performed with the KAPA SYBR FAST Universal Kit (Peqlab) and analyzed with the RotorGene RG-3000 (Corbett Research) using Ki67 primer (forward: TCT TGG CAC TCA CAG TCC AG; reverse: GCT GGA AGC AAG TGA AGT CC; Biolegio) and RPL13 as housekeeping gene (forward: TCT CCG AAA GCG GAT GAA CAC, reverse: CAA CAC CTT GAG GCG TTC CA; Biolegio). For BrdU experiments, 10 µM BrdU (Fluka) was added to the cell culture medium for 6 hours before fixation with 4% PFA for 10 minutes. For DNA denaturation, 2 M HCl was used for 30 min. Anti-BrdU antibody (mouse, clone BU-33, B8434, Sigma-Aldrich) was diluted 1:100.

### Immunocytochemistry

To analyze the differentiation in cell culture, Sox2 was used as a neuronal stem cell marker (mouse, MAB2018, R&D Systems, 1:100), Tuj1 (mouse, MAB1195, R&D Systems, 1:100) as a marker for early neurons, GFAP (rabbit, 180063, Invitrogen, 1:1000) for astrocytes, and CNPase (mouse, clone 11-5B, MAB326, Millipore, 1:5000) for labeling oligodendrocyte precursor cells. Fluorescein-labeled anti-mouse or anti-rabbit IgG (1:200, Invitrogen) was used as a secondary antibody. The HCN subtypes were detected using the following antibodies (1:100): HCN1: mouse, N70/28 (NeuroMab); HCN2: mouse, 75-111 (NeuroMab); HCN3: mouse, 75-175 (NeuroMab); HCN4: mouse, 75-150 (NeuroMab), followed by Alexa Fluor® 546 goat anti-mouse IgG (A-11030, 1:200, Molecular Probes).

For immunocytochemical stainings, cells were washed with PBS and fixed with 4% PFA for 10 minutes. After washing with PBS, cells were permeabilized and blocked in 0.1% TritonX-100 and 10% normal goat serum in PBS. The primary antibody, diluted to optimal working concentration, was applied overnight in 3% goat serum at 4°C. Cells were washed with PBS, followed by secondary antibody incubation (1 h at room temperature) with fluorescein-labeled anti-mouse IgG (1:200, Invitrogen) in 3% normal goat serum in PBS. Cells were counterstained by a 5-min incubation in Hoechst 33342 (Invitrogen) diluted 1:500. Slides were mounted using Fluoromount (Southern Biotech). All microscopic investigations were performed with an inverted fluorescent phase-contrast microscope (Keyence BZ-9000E). Ten randomly chosen images from each well were taken for analysis.

### Trans-well migration assay

Migration was analyzed via a trans-well migration assay using a modified Boyden chamber (CytoSelectTM 24-Well Cell Migration Assay, 8μM pore size, Cell Biolabs, Inc.) according to the manufacturer’s protocol.

### LDH assay

LDH release into the medium of the cultured cells was measured using the LDH Cytotoxicity Assay Kit (Thermo Scientific). Briefly, the medium (50 μl) was mixed with 50 μl of reaction mixture. After 30 min, 50 μl of stop solution was added, and the absorbance was read at 490 nm and 680 nm. To determine the LDH release, the 680-nm absorbance value was subtracted from the 490-nm absorbance.

### Live-/Dead-cell assay

The LIVE/DEAD® Cell-Mediated Cytotoxicity Kit (life technologies) was used to analyze the number of apoptotic cells. NSCs were stained with propidium iodide (1:500) as a non-permeable dye and simultaneously stained with Hoechst 33342 (1:500). After 5 minutes at 37°C, cells were counted using the inverted fluorescent phase-contrast microscope (Keyence BZ-9000E).

### Calcium imaging

Rat NSCs (NSC001, R&D systems, P#0-1) were plated on 35-mm dishes (µ-Dish 35mm, high Grid-500 Glass Bottom, Ibidi), and cultured according to the manufacturer’s protocol. Cells were loaded with 2.5 µM Fluo-4 AM (ThermoFisher) at 37°C for 15 min in complete medium and subsequently washed and incubated in imaging buffer (medium containing 10 mM HEPES, pH7.4) for 10 min. Imaging was performed on a TCS SP8 confocal microscope (Leica) with a 20X-immersion objective at 1024×1024 pixel resolution. Excitation was performed with WLL at 494 nm with 600-Hz scanning speed and 3s per frame. Baseline activity was recorded for 90 frames (270s). Subsequently, *I*_*h*_ blockage by ZD7288 treatment (Sigma-Aldrich) was assessed with 180 frames (540s), including 30 frames (90 s) of baseline recording, before manual addition of ZD7288 (30 µM) and subsequent recording of 120 frames (360 s) after ZD7288application, followed by final adjustment of external KCl to 40 mM for additional 20 recording frames (60s) to verify the viability of cells based on their ability to display a Ca^2+^signal after strong depolarization.

Imaging analysis was performed in ImageJ (78) with a custom-built macro domaindetect.ijm (according to (79)) for single-cell identification. After manual correction of over-segmentation and under-segmentation, each cell’s relative Ca^2+^signals were calculated from fluorescence intensities as ΔF / F_0_ for each time t, with F_0_ = mean baseline fluorescence intensity and ΔF = F_t_ - F_0_. Data are from n = 8 dishes (before and after treatment) with n = 777 vs. 676 total cells (38-138 cells per dish, mean ± SD = 90.8 ± 25.6).

### Electrophysiology

#### Two-electrode voltage-clamp experiments in *Xenopus laevis* oocytes

*Xenopus laevis* oocytes were obtained from tricaine-anesthetized animals. Ovaries were treated with collagenase (3 mg/ml, Sigma-Aldrich Chemie GmbH) in OR2 solution (NaCl 82.5 mM, KCl 2 mM, MgCl2 1 mM, HEPES 5 mM, pH 7.4) for 2-3 h, and subsequently stored in gentamycin solution (NaCl 75 mM, KCl 2 mM, CaCl_2_ 2 mM, MgCl_2_ 1 mM, HEPES 5 mM, pH 7.4, sodium pyruvate (550 mg/l), and gentamycin (50 mg/l)) at 18 °C. Oocytes were injected with cRNA encoding mHCN1 (0.5 ng), or HCN-DN (0.5 ng). The effect of HCN-DN on mHCN1 was verified by the co-expression of the two constructs with the pore mutant at a 1:1 ratio. Standard two-electrode voltage-clamp recordings were performed at room temperature (21–23 °C) three days after injection. ND66 solution (NaCl 66 mM, KCl 32 mM, CaCl_2_ 1.8 mM, MgCl_2_ 1 mM, HEPES 5 mM, pH 7.4) was used as a perfusion solution. The pipette solution contained 3 M KCl.

#### Electrophysiological recordings from adult mouse brain slices

For validation of the dominant-negative effect of the HCN-DN transgene on *I*_*h*_ in CA1 pyramidal neurons, 9- to 12-week-old B6-Tg (CaMKIIα-tTA^1Mmay/J^, (Bi-tetO-HCN^p.G480S^, eGFP)^C Isb^)) mice were used. Coronal hippocampal slices (300 µm) of adult mice were sectioned after intracardial perfusion with ice-cold sucrose–artificial cerebrospinal fluid (ACSF) (50 mM sucrose, 75 mM NaCl, 25 mM NaHCO_3_, 2.5 mM KCl, 1.25 mM NaH_2_PO_4_, 0.1 mM CaCl_2_, 6 mM MgCl_2_, and 2.5 mM glucose, oxygenated with 95% O2/ 5% CO2). After 60 min of recovery, slices were transferred to a recording chamber and continuously perfused at 2–4 ml min^−1^ with oxygenated ACSF (2.5 mM glucose, 22.5 mM sucrose) at 36°C. Fast excitatory and inhibitory synaptic transmission was inhibited by 20 µM CNQX (6-cyano-7-nitroquinoxaline-2,3-dione) and 10 µM gabazine. CA1 pyramidal neurons were visualized by infrared differential interference contrast (IR-DIC) video microscopy and epifluorescence for the detection of eGFP. Whole-cell patch recordings, data acquisition, and analysis were essentially performed as described (80).

#### Electrophysiological recordings from primary cortical stem cells

Neural stem cells were analyzed electrophysiologically using an Axiovert 200 (Zeiss) microscope and an EPC9 amplifier (HEKA). Pipets were pulled using a Sutter Puller Model P-1000 from borosilicate pipettes (GB150F-10, 0.86 × 150 x 100 mm; Science Products) with a pipette resistance between 2-6 MΩ. For voltage-clamp recordings, pipettes were filled with 10 mM NaCl, 130 mM KCl, 0.5 mM MgCl_2_, 5 mM HEPES, 1 mM EGTA, 3 mM Mg-ATP, 0.5 mM NaGTP (Abcam), pH 7.4. The bath solution contained 110 mM NaCl, 30 mM KCl, 5 mM HEPES, 1.8 mM CaCl_2_, 0.5 mM MgCl_2_, pH7.4, and was partially supplemented with 50 µM lamotrigine (Sigma). All cells analyzed exhibited a series resistance < 15 MΩ and were recorded using a 10-kHz Bessel filter. Data were recorded with Pulse software (HEKA) and analyzed with PulseFit or Fitmaster software (HEKA), respectively.

#### *In utero* electroporation

*In utero* electroporation was performed at E15 by exposing the uterus of the anesthetized dam and injecting 1-2 µl DNA (0.5-10 µg/µl) into the ventricle of the embryo. For visualization of the DNA in the ventricle, 0.05% Fast Green was added to the injection solution. Five pulses of 30-45 V were applied by the tweezertrodes (BTX, Harvard Apparatus) targeting the cortex. The electroporated construct pCAGIG-IRES-GFP (Addgene) was used as a control, and either the HCN-DN or the HCN-WT sequence was inserted into the experimental vector. The embryos were sacrificed at E19 for analysis.

#### Antibody validation

All antibodies used are commercially available and were selected for their specificity according to the manufacturer’s validation. For staining with each antibody, we additionally performed appropriate negative controls (omission of the primary antibody, or substitution by the normal serum from the animal in which the antibody was raised).

#### Statistical analyses

Statistical analyses were performed using GraphPad Prism software version 7.02. A single experiment (n) was defined as data points from one mouse or one well (10 images/well) for cell culture experiments. All data were tested for their normal distribution (D’Agostino and Pearson or Shapiro-Wilk normality tests) and, when appropriate, for variance of distribution (F test; Brown Forsythe). For single comparisons, an unpaired two-tailed t-test, or a Mann-Whitney test, was performed. All tests were two tailed. For multiple comparisons, a two-way analysis of variance (ANOVA), or one-way ANOVA, was employed. Post-hoc analysis with the honest Tukey’s significant difference test was subsequently performed when appropriate. For non-parametric testing, Kruskal-Wallis one-way ANOVA with post-hoc Dunn’s test was performed. For distribution analyses, Chi-square test was used. The sample size in each experiment was determined based on the previous publications using a similar experimental setting conforming to ethical guidelines for animal research. No randomization method was used. Unless specified otherwise, group measures are given as mean ± standard error of the mean (s.e.m.) with a significance annotation of * *p* < 0.05, ** *p* < 0.01, *** *p* < 0.001, **** *p* < 0.0001, and significance was set at *p* < 0.05.

## Supporting information

scRNA-seq differentially expressed genes

scRNA-seq cell-cycle marker genes

scRNA-seq marker genes

## ACKNOWLEDGMENTS

We thank Kathrin Sauter for technical assistance with mouse line generation and genotyping, Dr. Irm Hermans-Borgmeyer for transgenic services, Dr. Axel Neu (University Medical Center Hamburg-Eppendorf, UKE, Germany) for help with cell culture patch-clamp experiments, Dr. Jan Sedlacik (UKE) for assistance with the MRI experiments, Dr. Stefan Blaschke (University Hospital Cologne) for support with the stem cell culture experiments, Dr. Christoph Möhl (Image Data Analysis Facility, DZNE) for providing the domaindetect.ijm macro for Ca^2+^-imaging analysis, and Prof. Peter Nürnberg and his team for help with the scRNA-seq experiments. We acknowledge the support of animal facilities for excellent mouse care (Hamburg: H.Voss (ZMNH team); Cologne: Esther Mahabir-Brenner (CMMC team), Maria Guschlbauer (Medical Faculty team). We are grateful to the authors of La Manno *et al*. (27) and the Linnarsson lab (Karolinska Institute, Stockholm, Sweden) for making their dataset available to the public already at a pre-publication stage.

Furthermore, we thank Dr. Bina Santoro (Columbia University, New York, NY, USA), Prof. Gaia Tavosanis, and Prof. Paolo Salomoni (DZNE, Bonn, Germany), and M. Elisabeth Ross (Weill Cornell Medicine, New York, NY, USA) for helpful discussions and comments on the manuscript.

This work was supported by grants from the German Research Foundation (DFG, IS63/3-2 and IS63/10-1 (FOR 2715: *Epileptogenesis of genetic epilepsies*) to D.I.).

