## Supplemental Figures S1-S11 for "Developmental HCN channelopathy results in decreased neural progenitor proliferation and microcephaly in mice"

**This PDF file includes:**

Figures S1 to S11

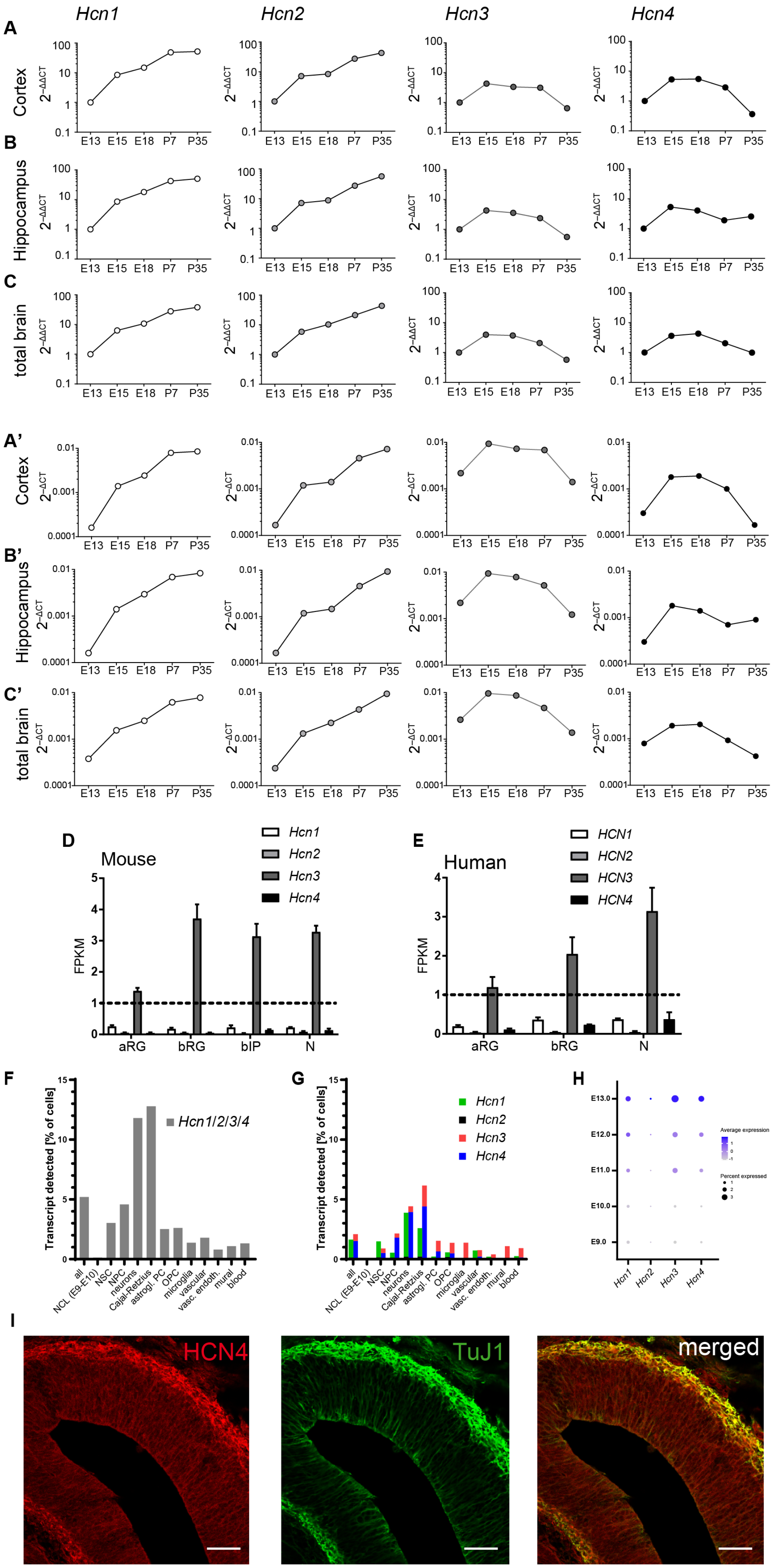
Hcn1
Hcn2
Hcn3
Hcn4
A
Cortex
B
Hippocampus
C
total brain
2−ΔΔCT
E13 E15 E18 P7 P35
A'
Cortex
B'
Hippocampus
C'
total brain
2−ΔCT
D
Mouse
FPKM
aRG
bRG
bIP
N
E
Human
HCN1
HCN2
HCN3
HCN4
F
Transcript detected [% of cells]
Hcn1/2/3/4
all
NCL (E9-E10)
NSC
NPC
neurons
Cajal-Retzius
astrogl. PC
OPC
microglia
vascular
vasc. endoth.
mural
blood
G
H
E13.0
E12.0
E11.0
E10.0
E9.0
Average expression
Percent expressed
I
HCN4
TuJ1
merged

**Figure S1: Endogenous HCN-subtype expression in mouse, rat, and human tissues.**

The mRNA expression levels of the *Hcn1-Hcn4* subtypes in mouse **(A)** cortex, **(B)** hippocampus, and **(C)** total brain normalized to *GAPDH* and the corresponding E13 time point shown as the $2^{-\Delta\Delta CT}$ and the same regions normalized to GAPDH (A' – B') shown as the $2^{-\Delta CT}$ values. **(D)** Analysis of mRNA expression of HCN subtypes in apical radial glia (aRG), basal radial glia (bRG), basal intermediate precursor (bIP) cells, and neurons (N) in mouse and **(E)** aRG, bRG, and N in humans revealed the predominant presence of the HCN3 subunit in all cell types analyzed. **(F)** Single-cell RNA-seq analysis of a publicly available data set from the Linnarsson lab (27) combines E9, 10, 11, 12, 13 mouse brains and showed mRNA expression of all HCN subunits in almost all identified cell types, with **(G)** HCN4 and HCN3 as the most abundant subunits in neuronal progenitors. **(H)** The overall expression of HCN1, HCN3 and HCN4 is increasing during the represented embryonic period (E9 – E13) in expression levels and cell numbers. **(I)** Immunofluorescence staining of coronal section of the mouse cortex at E13 revealed expression of HCN4 (red) in young neurons positive for TuJ1 (green). Scale bars 50 µm; Data are presented as mean ± s.e.m; (A, B, C,) n=1, (D) n=4, (E) n=3/2; Images in (I) are representative of three similar experiments.

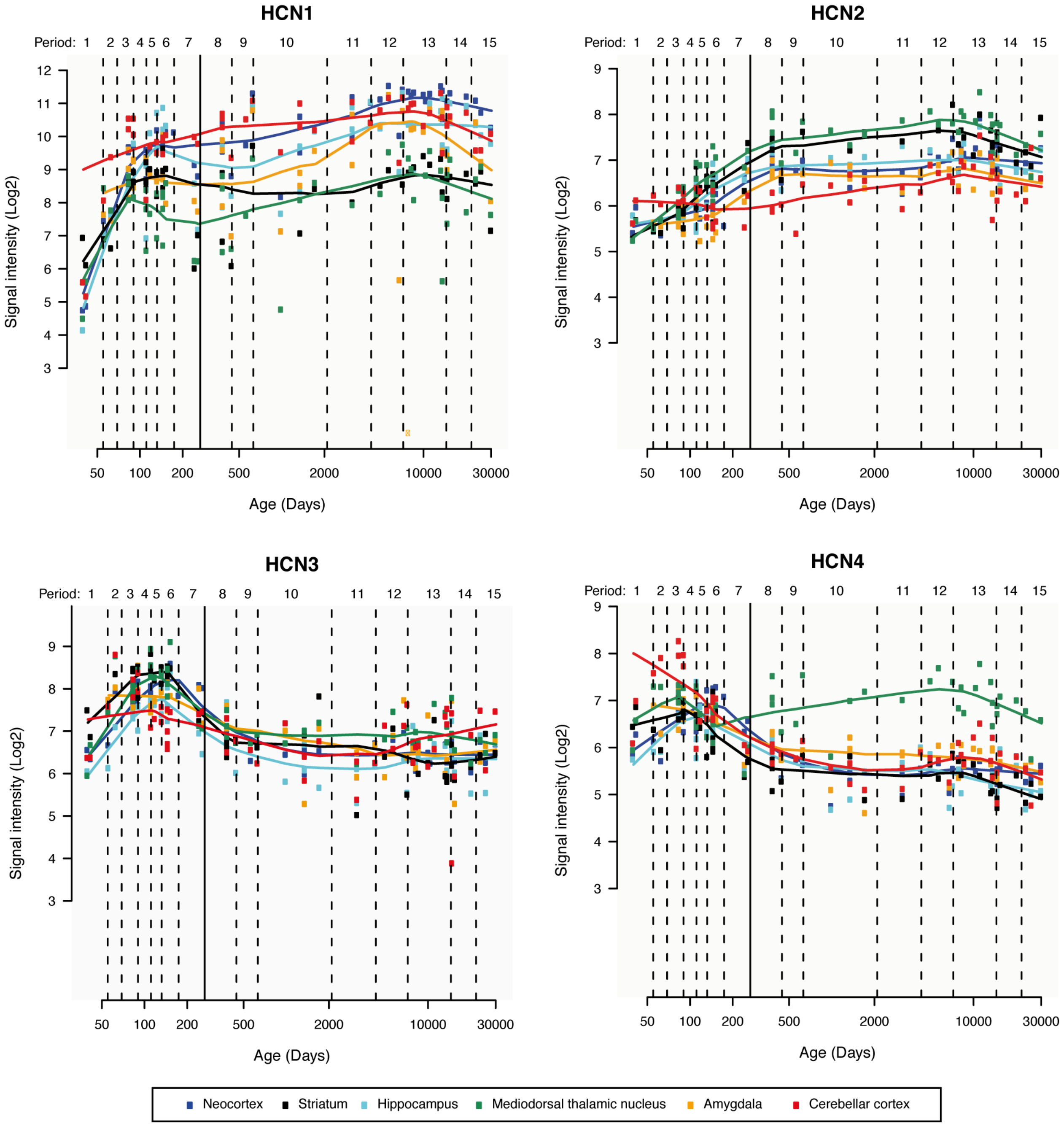


**Figure S2: Spatio-temporal expression of HCN subunit transcripts in the human brain.**

The expression levels of the HCN1-4 transcripts during human brain development were obtained from the online dataset of Kang *et al.*, 2011 (27) available at http://hbatlas.org/pages/hbtd.

Age is given in post-conceptional days (late embryonic development (4-8 weeks post conception, period 1), fetal development (periods 2–7), postnatal development (periods 8–12), and adulthood (periods 13–15)), the gray vertical line indicates the time point of full-term birth, expression levels are given as array signal intensity (25). The brain regions are listed in the legend.

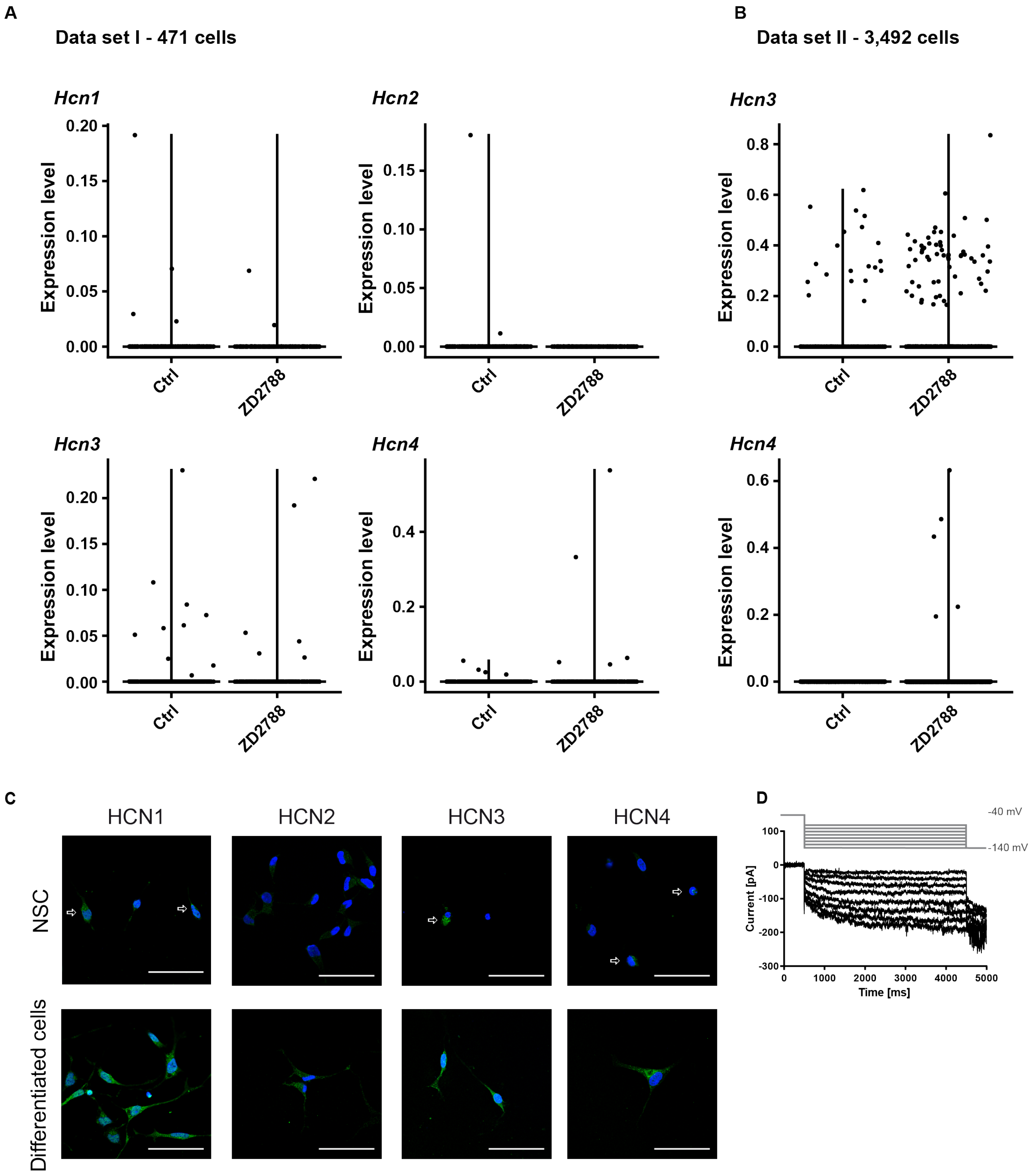


**Figure S3: Expression levels of the four HCN subtypes in two distinct single cell RNA sequencing data sets as well as their protein expression and functionality in rat neuronal stem cells.**

**(A)** In the WaferGen scRNAseq dataset (I) of the 471 cells analyzed 7 were expressing Hcn1 3 Hcn2, 20 Hcn3 and 9 cells Hcn4. The combination of Hcn3+Hcn4 and Hcn2+Hcn3 was found each in 1 cell. **(B)** Using the 10x Genomics platform a total of 3,492 cells could be analyzed, here only 80 expressed HCN3, 5 expressed HCN4, and only 1 cell showed HCN3+HCN4 expression. **(C)** Immunofluorescence staining of cultured primary cortical stem cells revealing HCN1, HCN3, and HCN4 expression (arrows), but no detectable HCN2 immunoreactivity, while in differentiated cells from the same rat cell culture all HCN subtypes were detectable. **(D)** Two out of 25 neural stem cells analyzed showed an $I_h$-like current upon hyperpolarization to –140 mV for 4 s. The cell was recorded in a bath solution supplemented with 50 µM lamotrigine. Scale bars 50 µm. Images in (C) are representative of four similar experiments.

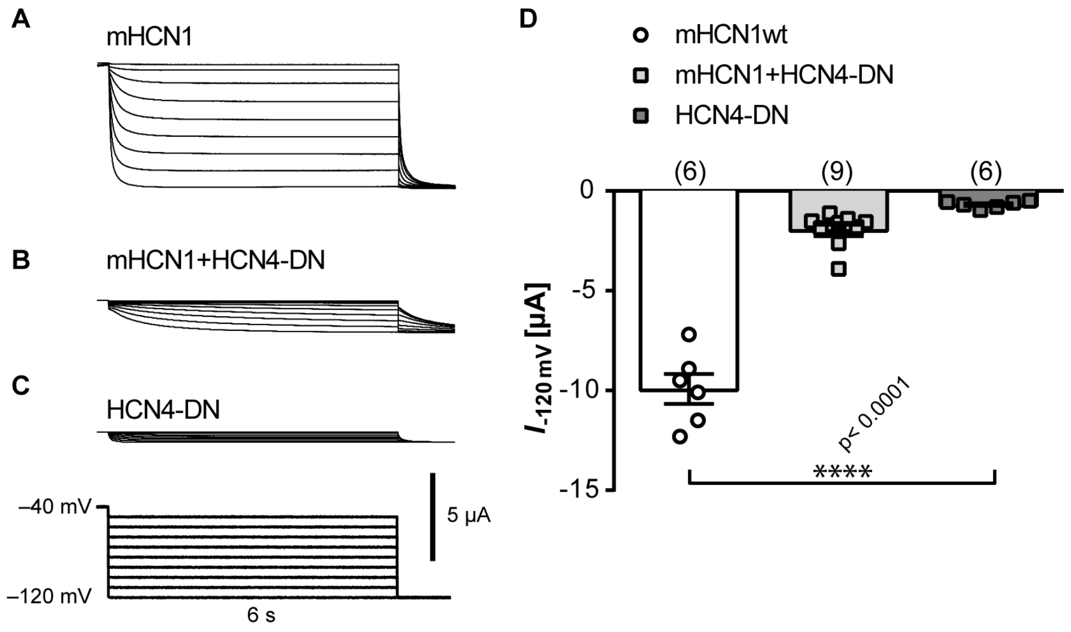


**Figure S4: HCN-DN is a dominant-negative subunit suppressing mHCN1-mediated currents.**

Representative traces of two-electrode voltage-clamp recordings in *Xenopus laevis* oocytes injected with equimolar ratios of complementary RNA encoding (**A**) mouse wild-type HCN1 (mHCN1), (**B**) mHCN1 and the dominant-negative HCN4 subunit (HCN-DN), and (**C**) HCN4-DN. The voltage-clamp protocol is shown below the traces. (**D**) Measurements at –120 mV demonstrated a dominant-negative effect of HCN-DN on mHCN1-mediated currents (Kruskal-Wallis with post-hoc Dunn's test). Data are presented as mean ± s.e.m; n is given in parentheses; p values are above asterisks.

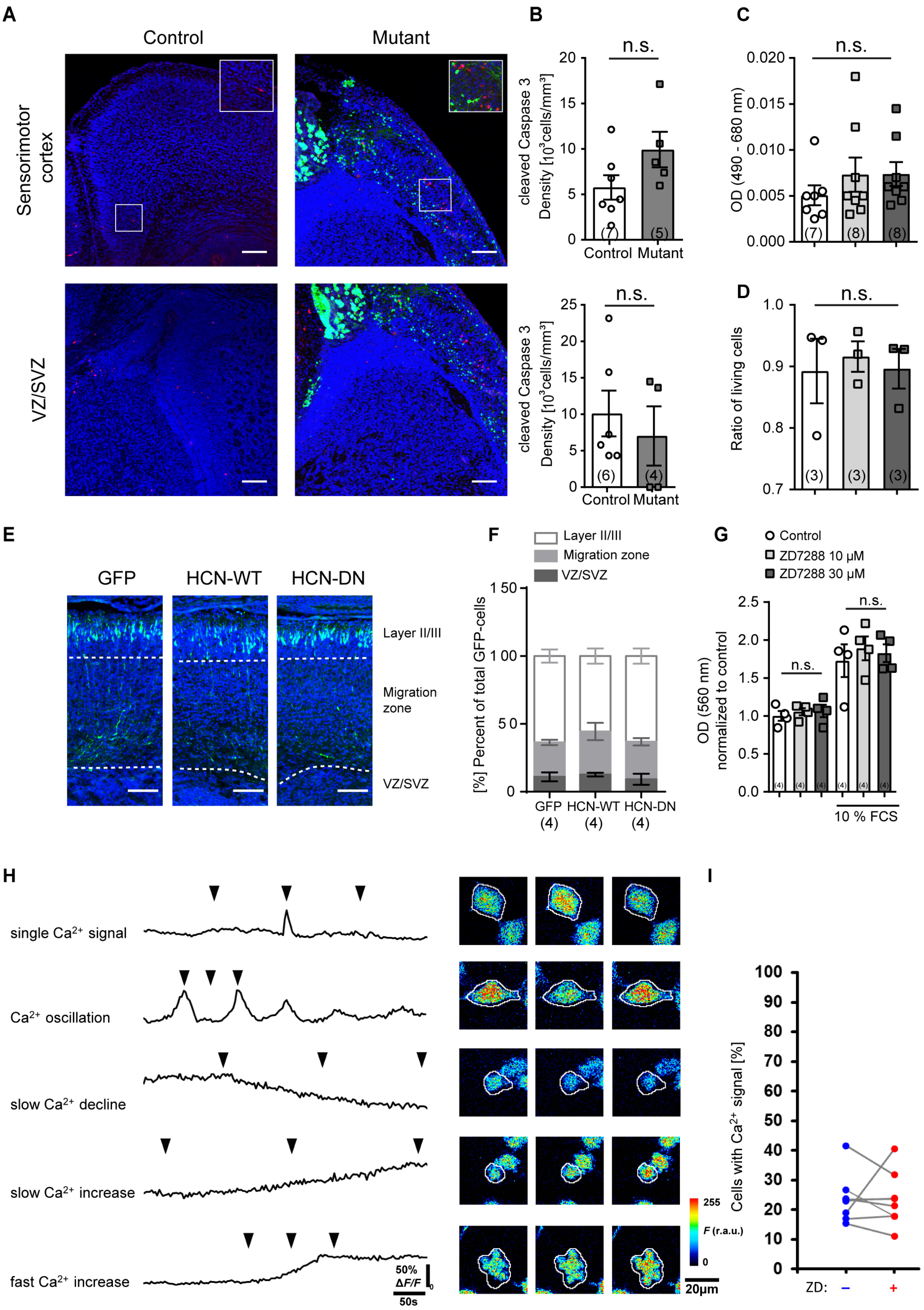


**Figure S5: Loss of $I_h$ function effects on apoptosis, migration, or intracellular calcium handling.**

**(A)** Representative images of the sensorimotor cortex and VZ/SVZ of neonatal (P0) control and mutant EMX1-HCN-DN mice stained against the apoptosis marker *cleaved Caspase-3*.

**(B)** Unlike in E12.4 embryos, quantification of the *cleaved Caspase-3* staining in P0 brain slices revealed no differences in the number of apoptotic cells in sensorimotor cortex or VZ/SVZ (Mann-Whitney test). **(C)** Analysis of LDH activity as a marker of cell damage revealed no apoptotic or necrotic effect of 10 µM or 30 µM ZD7288 application on cortical stem cells (Kruskal-Wallis). **(D)** Live/dead assay showed no effect of ZD7288 (10 µM and 30 µM) on apoptosis of the primary rat cortical stem cells (Kruskal-Wallis). **(E)** Representative images of *in utero* electroporation at E15 showed no difference in neuronal migration at E19. **(F)** Quantification of migration depicted as percentage of GFP-positive cells [%] in layer II/III, migration zone, including layers IV to VI, intermediate zone and cortical plate, and VZ/SVZ showed no significant difference (two-way ANOVA). **(G)** Trans-well migration assay of primary cortical stem cells of the rat indicated no differences in the migration of ZD7288-treated vs. non-treated cells at baseline (left bars), nor did it show differences after addition of the chemoattractant fetal calve serum (FCS) (optical density (OD) measured at 560 nm, two-way ANOVA with post-hoc Tukey's test). **(H)** Representative $Ca^{2+}$-signals in primary rat cortical stem cells, fluorescence intensity-colored pictures (*right*, signal ROI marked with white border) correspond to time points indicated by arrows above respective traces (*left*). **(I)** No difference could be detected in the total percentage of cells displaying $Ca^{2+}$-signals before or after $I_h$ blockage by ZD7288 (23.7 ± 2.9 % vs. 23.5 ± 3.2 %, $p > 0.95$ with paired t-test; n = 777 / 676 cells from n = 8 culture dishes). Scale Bars 100 µm; Data are presented as mean ± s.e.m; n is given in parentheses; Images in (A) are representative examples of (B); Images in (E) are representative examples of (F).

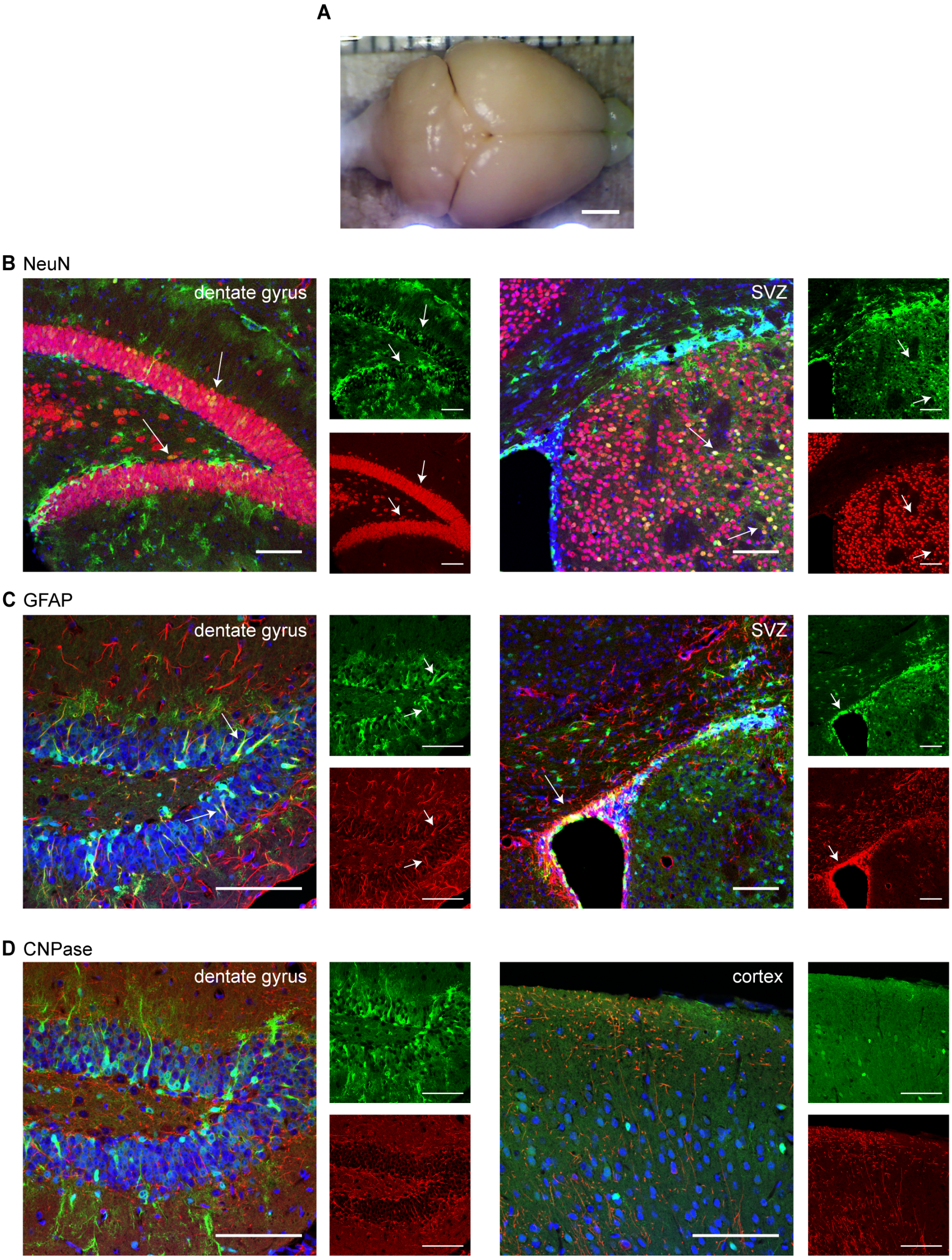


**Figure S6: Deletion of the C-terminus of the HCN channel does not change gross morphology in adult EMX1-HCN4-573X.**

**(A)** Representative image of the total brain of an adult EMX1-HCN4-573X mouse shows no gross morphological alteration. Immunostaining against **(B)** the neuronal marker NeuN (red_, **(C)** the astrocyte marker GFAP (red) or **(D)** the oligodendrocyte marker CNPase (red) show no structural changes in the dentate gyrus (right) and SVZ or cortex (left) of EMX1-HCN4-573X adult mice, eGFP is depicted in green, DAPI shown in blue. Scale Bars in (A) 2 mm, (B - D) 100 µm; Images are representative examples of two experiments.

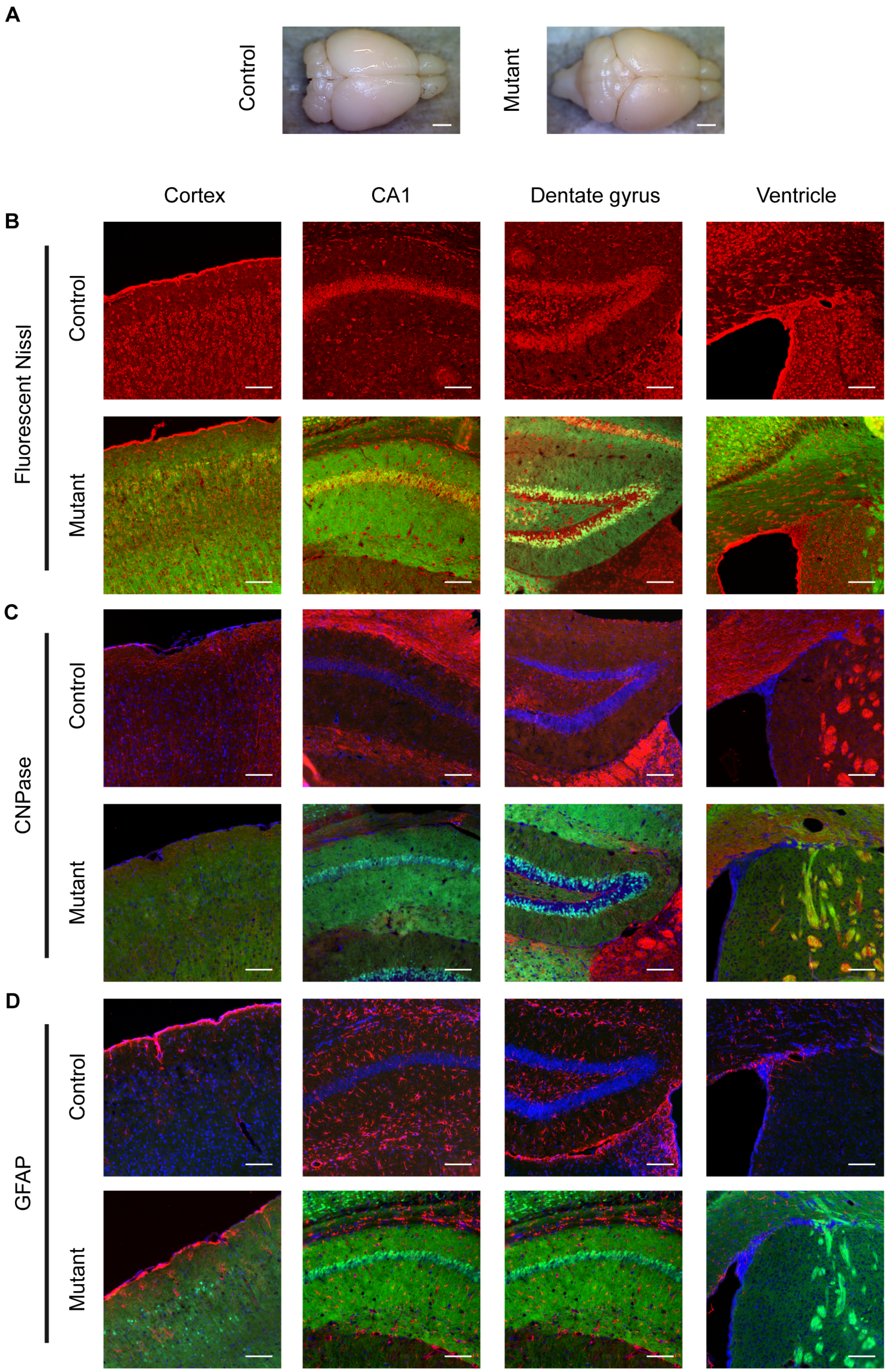


**Figure S7: Forebrain structure is unaffected in NEX-HCN-DN mutants.**

**(A)** Explanted brains of adult NEX-HCN-DN mice expressing the HCN-DN transgene under the control of the NEX promoter, which starts expression at E10.5 in mainly post-mitotic forebrain progenitors and neurons, did not exhibit gross morphological changes as compared to control mice. **(B)** In coronal sections, staining with fluorescent Nissl dye (red) showed no gross morphological differences in the cortex, hippocampus (CA1 and dentate gyrus), or the VZ/SVZ between NEX-HCN-DN mutants and controls. Example images of the cortex, hippocampus (CA1 and dentate gyrus), and the VZ/SVZ showed no differences in the staining patterns for **(C)** oligodendrocytes (CNPase, red) and **(D)** astrocytes (GFAP, red). eGFP is depicted in green, DAPI is shown in blue. Scale bars in (A) 2 mm, (B – D) 100 µm; Images in (A) are representative of three similar experiments, in (B –D) images are representative of two similar experiments.

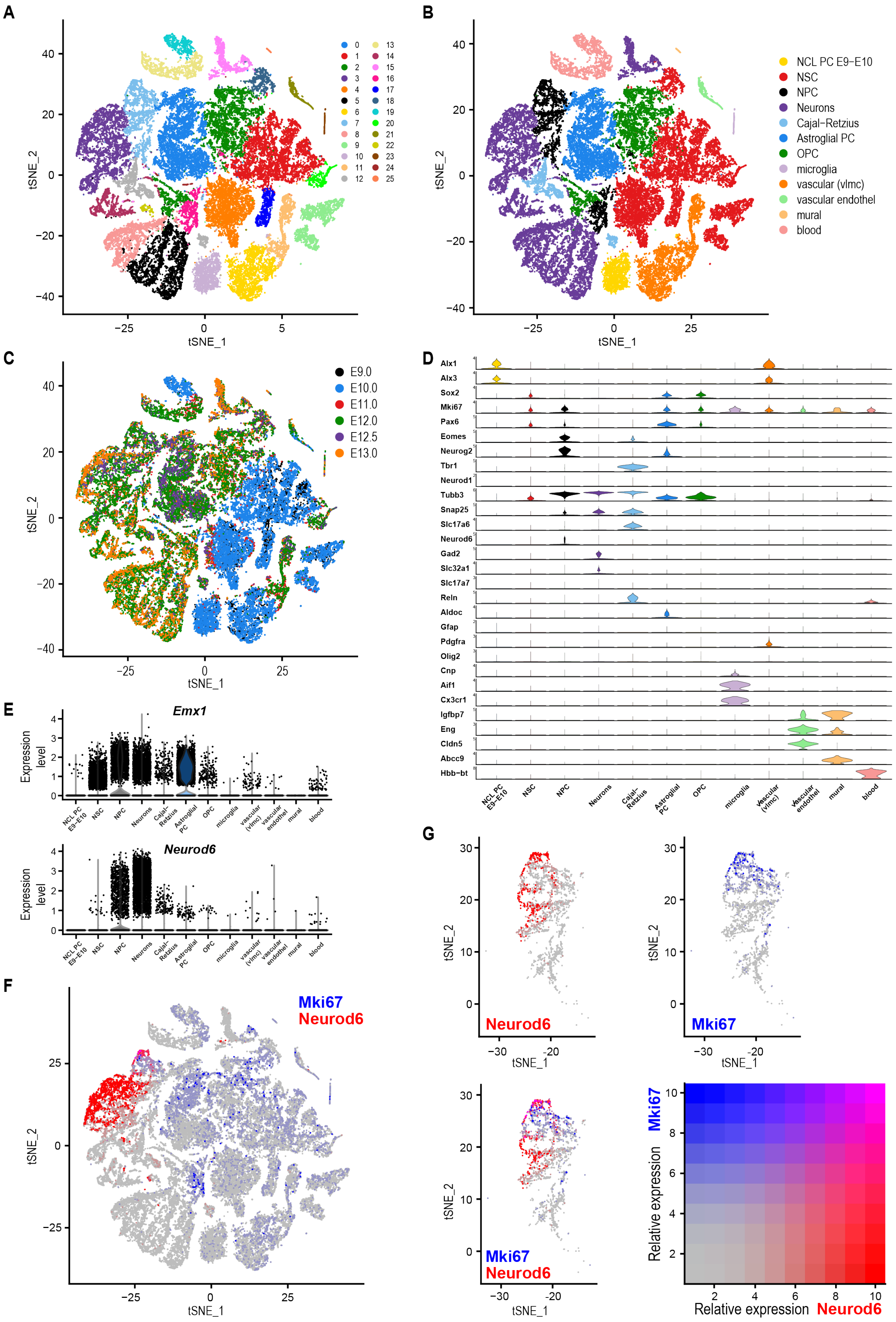
A
B
NCL PC E9−E10
NSC
NPC
Neurons
Cajal−Retzius
Astroglial PC
OPC
microglia
vascular (vlmc)
vascular endothel
mural
blood
C
E9.0
E10.0
E11.0
E12.0
E12.5
E13.0
D
Alx1
Alx3
Sox2
Mki67
Pax6
Eomes
Neurog2
Tbr1
Neurod1
Tubb3
Snap25
Slc17a6
Neurod6
Gad2
Slc32a1
Slc17a7
Reln
Aldoc
Gfap
Pdgfra
Olig2
Cnp
Aif1
Cx3cr1
Igfbp7
Eng
Cldn5
Abcc9
Hbb−bt
E
Emx1
Neurod6
Expression level
F
Mki67
Neurod6
G
Relative expression
tSNE_1
tSNE_2

**Figure S8: Single-cell RNASeq analysis of a publicly available data set from the Linnarsson lab (27) combines E9, 10, 11, 12, 13 mouse brains (data set III).**

**(A)** The scRNAseq data set clustered in 26 distinct clusters, which can be classified in **(B)** 12 different cell types. **(C)** The tSNE projection shows the embryonic time points of the analyzed tissue. **(D)** Violin plots depict the expression level of the maker genes of each identified cell type. **(E)** Expression of *Emx1* is detectable in NSCs, as well as in the different progenitor cells and neurons, while *Neurod6* (NEX) expression is restricted to NPCs and neurons in the different cell types. **(F-G)** Throughout the analyzed time points the number of cells co-expressing *Neurod6* (red) and *Mki67* (blue) is low shown in the whole data set (F) as well as in the NPC (G), only cluster 7 is shown, because cluster 16, did not express any *Neurod6*.

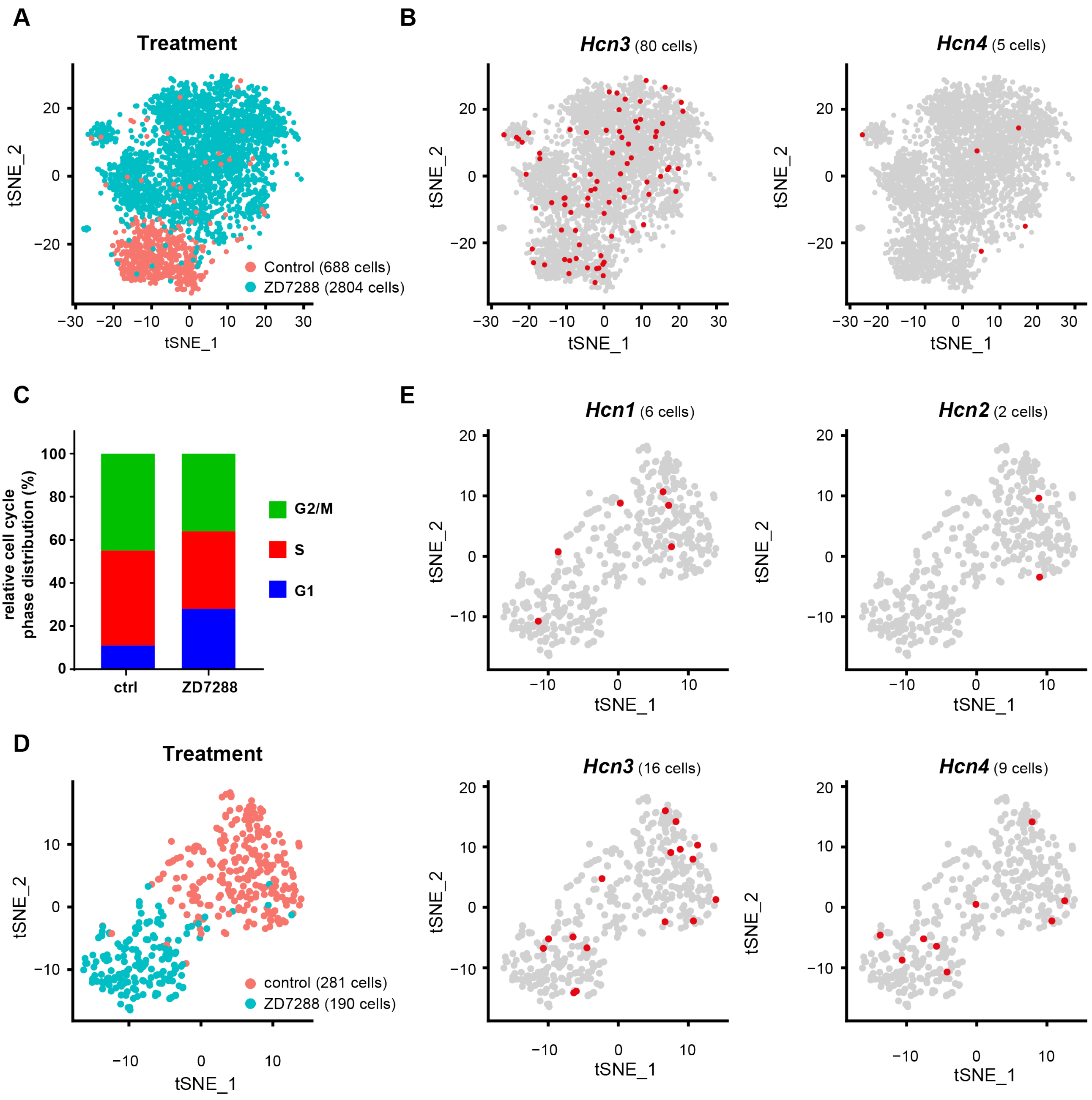


**Figure S9: Single-cell RNA sequencing Data obtained using 10x Genomics platform (data set II) shows no discriminate clustering of HCN expression cells, similar to the Wafergen data set (I).**

**(A)** t-SNE projection of scRNA-seq data of untreated (Control, salmon, 688 cells) and ZD7288-treated (30 µM) cells (Treatment, green, 2804 cells)). **(B)** HCN3 and HCN4 expressing cells depicted on the tSNE projection of dataset II. **(C)** Similar to the results of data set I (see Fig. 3), the percentage of cells in the G1, S, or G2/M cell cycle phases showed a significant difference in the distribution of ZD7288 versus control (vehicle-treated) cells ($p < 0.0001$, Chi square). **(D)** In data set I 281 untreated, control cells (salmon) and 190 ZD7288-treated cells were analyzed. **(E)** Cells expressing the HCN1 – 4 subunits in data set I shown in tSNE projections

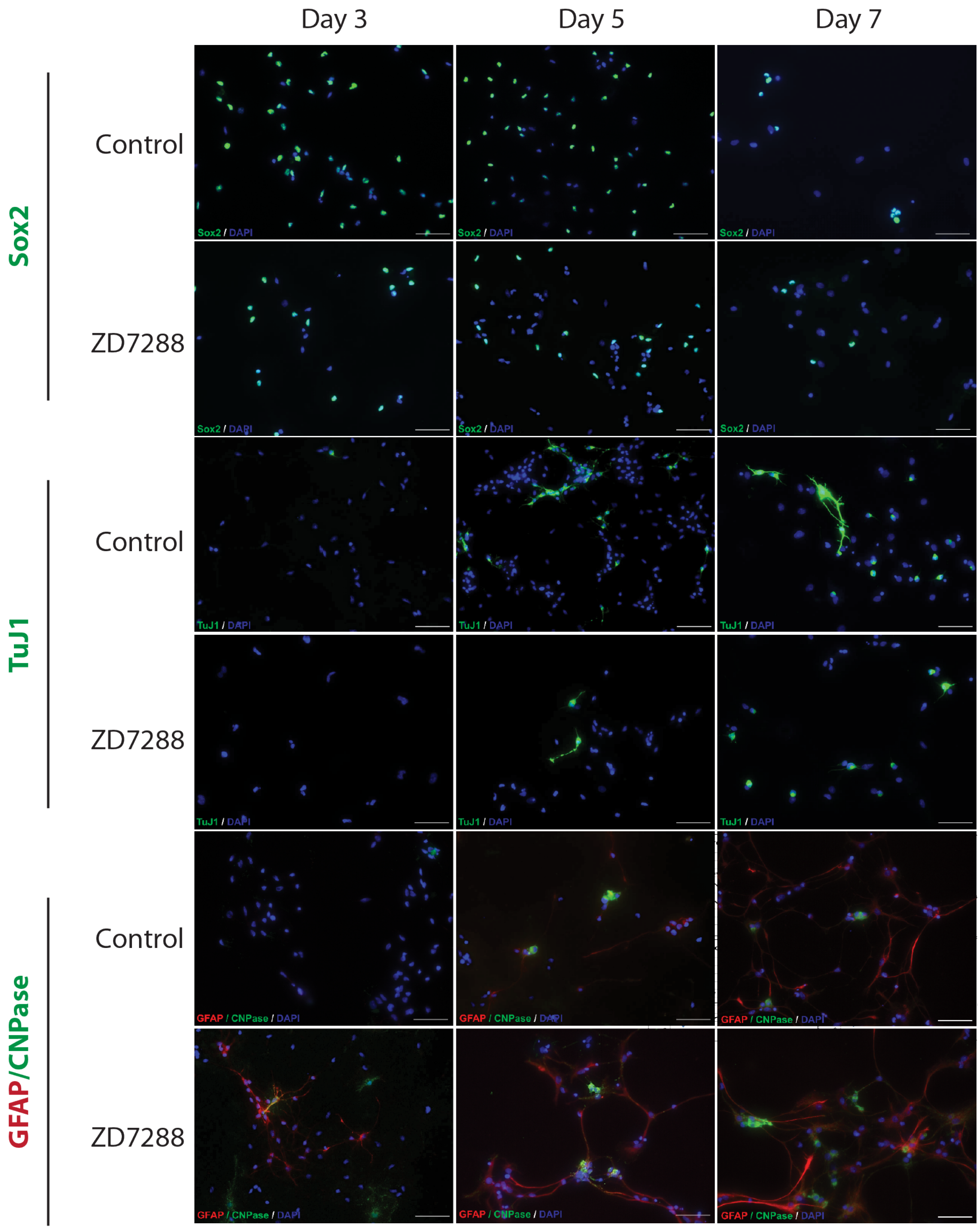


**Figure S10: Differentiation of control and 30 µM ZD7288 treated rat NSCs *in vitro*.**

Representative immunofluorescence images of control rat NSC or NSCs treated with 30 µM ZD7288 on day 3, 5 and 7 after differentiation upon FGF2 withdrawal. Cells were stained for neural progenitors (Sox2 in green; first two rows), early Neurons (TuJ1 in green; third and fourth row) as well as for astrocytes (GFAP in red) and oligodendrocytes (CNPase in green, last two rows). Scale bar 50 µm.

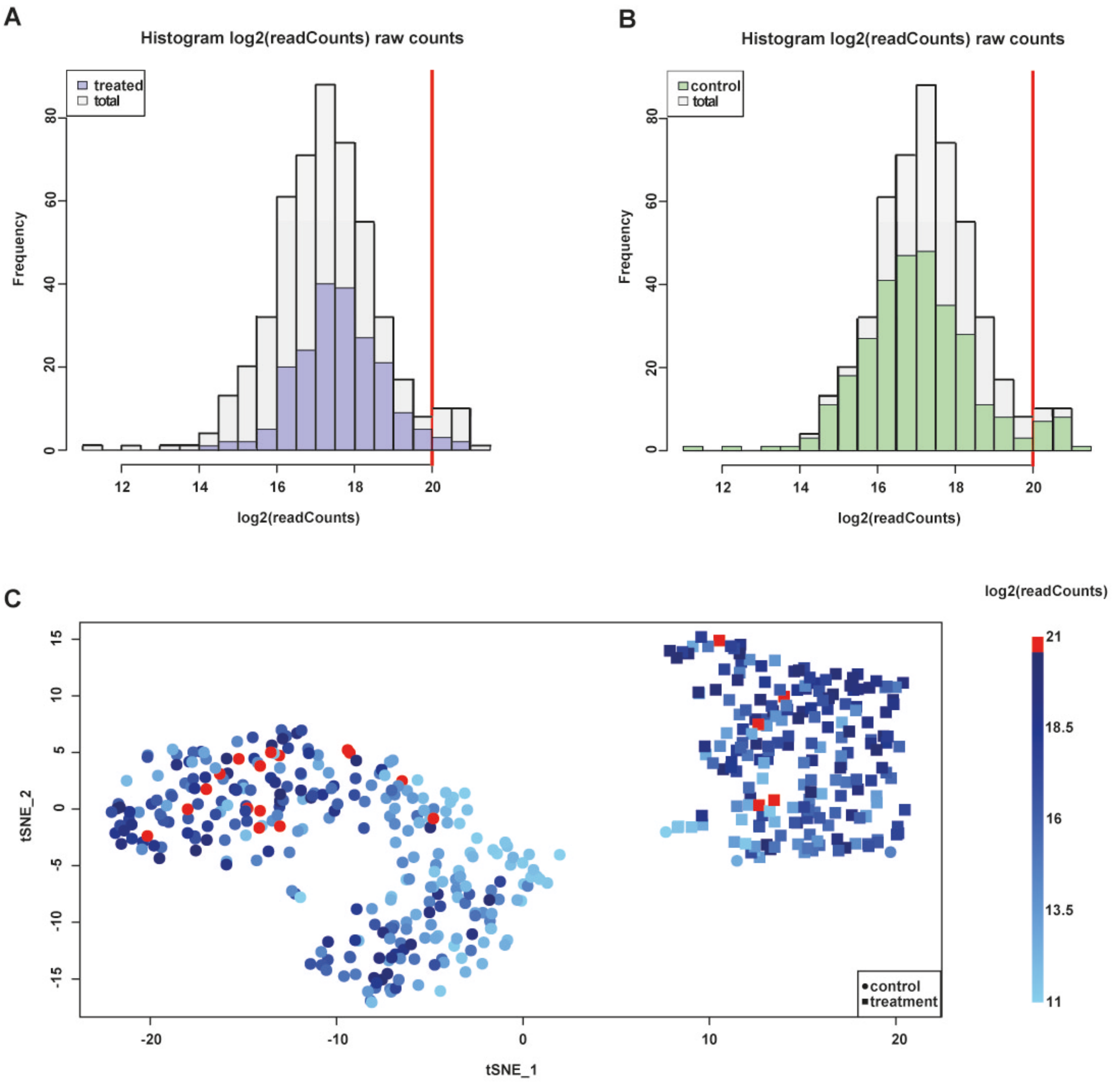


**Figure S11: Distribution of scRNA-seq read coverage.**

**(A, B)** Distribution of mapped reads ($\log_2$ counts) per barcode, for all barcodes (grey bars), and for barcodes corresponding to ZD7288 treated cells (blue bars) and untreated cells (green bars). The vertical line at 220 counts marks the boundary above which the counts are likely to originate from multiple cells. Those barcodes were excluded from further analysis. **(C)** 2-dimensional t-SNE plot of all barcodes (according to their expression profile). Circles (rectangles) represent barcodes related to ZD7288-treated (non-treated) cells. Colors correspond to the total number of read counts per barcode. Excluded cells with more than $2^{20}$ counts are highlighted in red.
