## Supplementary material for "Developmental HCN channelopathy results in decreased neural progenitor proliferation and microcephaly in mice": scRNA-seq marker genes

| Cluster | primary cell type | primary marker | specific marker |
|---|---|---|---|
| **neuron glutamatergic** | neuron | *Tubb3 (Tuj1), Snap25, Rab3a, Rbfox3* | *Slc17a7, Neurod6* |
| **neurons GABA-ergic** | neuron | *Tubb3 (Tuj1), Snap25, Rab3a, Rbfox3* | *Gad1, Gad2, Slc32a1* |
| **Cajal-Retzius cell** | neuron | *Tubb3 (Tuj1), Snap25, Rab3a, Rbfox3* | *Slc17a6, Reln* |
| **immature neurons / postmitotic neuronal progenitors** | progenitor | *Pax6* | *Eomes (Tbr2), Neurod1, Tbr1* |
| **neuronal stem cell (NSC)** | progenitor | *Pax6* | *Sox2, Mki67* |
| **oligodendrocyte precursor cell** | progenitor | *Sox10* | *Olig2, Pdgfra* |
| **oligodendrocyte** | glia | *Sox10* | *Cnp, Olig2* |
| **astrocyte** | glia | *Gfap* | *Aldoc, Aqp4* |
| **microglia** | immune | *Dock2* | *Aif1 (Iba1), Cx3cr1, Ly86* |
| **vascular endothel** | vascular | *Igfbp7, Eng* | *Ly6c1, Cldn5, Kdr* |
| **mural cell** | vascular | *Igfbp7, Eng* | *Rgs5, Abcc9, Pdgfrb* |
| **blood cell** | blood | *hemoglobins* | *Hbb-a1, Hbb-bs, Hbb-bt* |
| **neural crest E9-E10 precursor** | progenitor | | *Alx1, Alx3* |
